# Autoregulation of Many Yeast Ribosomal Proteins Discovered by Efficient Search for Feedback Regulation

**DOI:** 10.1101/2020.05.11.089300

**Authors:** Basab Roy, David Granas, Fredrick Bragg, Jonathan A. Y. Cher, Michael A. White, Gary D. Stormo

## Abstract

Post-transcriptional autoregulation of gene expression is common in bacterial systems but many fewer examples are known in eukaryotes. We used the yeast collection of genes fused to GFP as a rapid screen for examples of feedback regulation in ribosomal proteins by overexpressing a non-regulatable version of a gene and observing the effects on the expression of the GFP-fused version. We tested 95 ribosomal protein genes and found that 21 of them showed at least a three-fold repression. Two genes, *RPS22B* and *RPL1B*, showed over a 10-fold repression. In both cases the cis-regulatory segment resides in the 5’ UTR of the gene as shown by placing that segment of the mRNA upstream of GFP alone and demonstrating it is sufficient to cause repression of GFP when the protein is over-expressed. Further analyses showed that the intron in the 5’ UTR of *RPS22B* is required for regulation, presumably because the protein inhibits splicing that is necessary for translation. The 5’ UTR of *RPL1B* contains a sequence and structure motif that is conserved in the binding sites of Rpl1 orthologs from bacteria to mammals, and mutations within the motif eliminate repression.

## Introduction

Feedback regulation is common in many biological processes. In metabolic pathways the end product can often inhibit one of the enzymes in the pathway to set the flux through the pathway at the appropriate level^1,2^. Gene expression is also often regulated by feedback mechanisms. This commonly occurs with a transcription factor that regulates its own expression, referred to as autoregulation. One of the first transcription factors to be studied in detail, the lambda repressor, was found to be autoregulated both positively and negatively, allowing it to maintain its *in vivo* concentration in a narrow range^3,4^. Autoregulation has been found to be among the most common network motifs in bacterial transcription^5,6^. Studies on the regulatory network in yeast also identify many examples of autoregulation ^7–9^. Mathematical analyses have characterized the properties and advantages of autoregulatory networks ^10–12^.

Although less well-studied, autoregulation also occurs for proteins involved in post-transcriptional steps of gene expression. For example, many splicing factors regulate their own expression ^13–17^. Recently developed methods for high-throughput analysis of RNA-protein interactions have identified many RNA-binding proteins, some of which are associated with their own mRNAs ^18–25^. By itself that doesn’t prove they are autoregulatory, but it seems likely to be a consequence of such binding. In most of those cases, both transcription factors and RNA-binding proteins, the normal function of the protein is to bind DNA or RNA and often to regulate gene expression. The fact that they can regulate their own expression is not surprising given the advantages of such feedback processes, it only requires that the gene’s own DNA or RNA be included in the target list for the protein. Binding sites are often short and, because unconstrained nucleic acids can evolve rapidly, sites that offer a selective advantage are likely to be obtained through random mutagenesis processes.

There are also examples of proteins whose primary function is not in gene regulation but have a secondary role in regulating their own expression. Many of these are proteins that bind to RNA, but whose primary functions are not involved in controlling gene expression. For example, most of the ribosomal proteins in *E. coli* are subject to feedback regulation ^26,27^. Ribosomal proteins are expressed as part of transcription units (operons) composed of other ribosomal proteins. Autoregulation by one of the proteins in the operon is typically sufficient to control the expression of all the other genes in the operon by translational coupling, where translation of an upstream gene in an operon is required for translation of the downstream genes. The ribosomal proteins are all RNA-binding proteins, having as their primary target the rRNAs of the ribosome. To become autoregulatory, the mRNA simply has to evolve a sequence that is a molecular mimic of the primary target site, but with lower affinity so that binding of the rRNA is saturated before the regulatory site becomes bound by the protein ^27–30^. There are also examples of tRNA synthetase genes in bacteria and yeast that have evolved a similar regulatory site, where the mRNA mimics the tRNA that the synthetase gene normally binds to, but with lower affinity ^31–35^. Particularly interesting are cases where the protein’s normal function does not involve binding to RNA but it is found in screens for RNA-binding proteins ^36,37^. In some cases, proteins with alternative primary functions have been shown to be direct regulators of their own translation ^38–42^. Such examples highlight the enormous functional capacity of RNA where it can become a sensor of the cellular environment and autonomously regulate its own fate, as exemplified in cases where no protein is required, such as riboswitches ^43–47^. When the effector being recognized by the mRNA is its own gene product, the result is autoregulation of gene expression.

The fundamental characteristic of feedback regulation of gene expression is that if the activity of a gene product, which is usually proportional to its concentration, is higher than the set point, or “desired” level for the cell, then its expression is reduced, and conversely, if the activity is too low, the expression is increased. This relationship is true regardless of the mechanism by which the feedback regulation occurs, whether it involves a complex network of interactions or is simply the result of direct autoregulation by the gene product itself. Once examples of feedback regulation of gene expression are obtained, the mechanism can be determined by additional experiments. The collection of yeast strains with genes fused to green fluorescent protein (GFP) ^48^ provides an excellent resource to screen for examples of feedback regulation. By introducing into those strains an inducible copy of a gene for the same protein, but lacking all potential <u>c</u>is-<u>r</u>egulatory <u>e</u>lements (cre-less), an observed reduction in the level of GFP after induction indicates some feedback mechanism controlling the expression of the wild-type gene. Further analyses are required to determine the step in the expression process that is being regulated, whether it is transcription initiation, any of the processes leading to the mature mRNA, any step in translation, or even post-translational enhancement of protein degradation. We are particularly interested in finding new examples of post-transcriptional regulation of protein expression so our initial focus is on ribosomal proteins, which are commonly translationally autoregulated in bacteria but^27,28,49,50^ for which many fewer examples are known in yeast.

Ribosome synthesis in yeast is subject to feedback regulation in part by alternative functions of ribosomal proteins ^51–55^. There are several examples of post-transcriptional autoregulation by yeast ribosomal proteins, most often through inhibition of splicing necessary for protein expression. For example, *RPL22B, RPS14B* and *RPL30* all have introns within the N-terminus of the coding sequence and splicing is inhibited by binding of the encoded protein ^56–59^. Remarkably, the ortholog of Rpl30 in the archaeon *Sulfolobus acidocarldarius* can bind to the same mRNA target and inhibit splicing ^60^. *RPS9A* and *RPS9B* both have introns within the N-terminus of the coding region and both genes are subject to feedback regulation by inhibition of splicing ^61^. The orthologs of Rps9 are involved in autoregulation in several other eukaryotic species and even in bacteria ^27,61^. *RPS28B* does not contain an intron but is autoregulated by a different mechanism where binding of the Edc3 decapping enzyme to the 3’ UTR is regulated by the Rps28 protein, leading to mRNA degradation ^62,63^. These cases are all consistent with examples from bacteria where ribosomal protein synthesis is regulated post-transcriptionally, and it seems likely that a directed search for feedback regulation among yeast ribosomal protein genes could uncover more examples, leading us to utilize the yeast GFP-fusion collection.

## Results and Discussion

Feedback regulation of protein expression requires that when the activity of the protein, usually proportional to its concentration, is higher than the homeostasis point of the cell, its expression is reduced, and when the activity is lower than that point, its expression is increased. This allows the cell to maintain expression in a narrow range around its set point. The collection of yeast genes fused to GFP ^48^ provides an excellent resource to screen for genes that exhibit feedback regulation. A version of the gene is synthesized that is lacking any potential cis-regulatory elements (a cre-less version of the gene), under the control of an inducible promoter. The cre-less version is synthesized with alternative 5’ and 3’ UTRs, mCherry is fused to the C-terminus in place of the GFP of the wild-type gene, any introns are removed, and the codons of the gene are shuffled ^64^ to maintain the wild-type protein sequence while altering the mRNA sequence sufficiently that we expect any cis-regulatory elements that overlap the coding sequence would be eliminated. We use the *GAL1* promoter for induction of the cre-less gene (Figure 1a). If there is feedback regulation, overexpression of the cre-less gene (monitored by mCherry fluorescence) will bind to the regulatory elements of the wild-type gene, leading to a decrease in its expression, which is monitored by GFP fluorescence. Identification of feedback regulated genes does not provide information about the mechanism of action, and further analyses are required to determine the step in the expression process that is regulated. The scheme for gene synthesis and fluorescent detection is summarized in supplementary figure S1.

**Figure 1:**
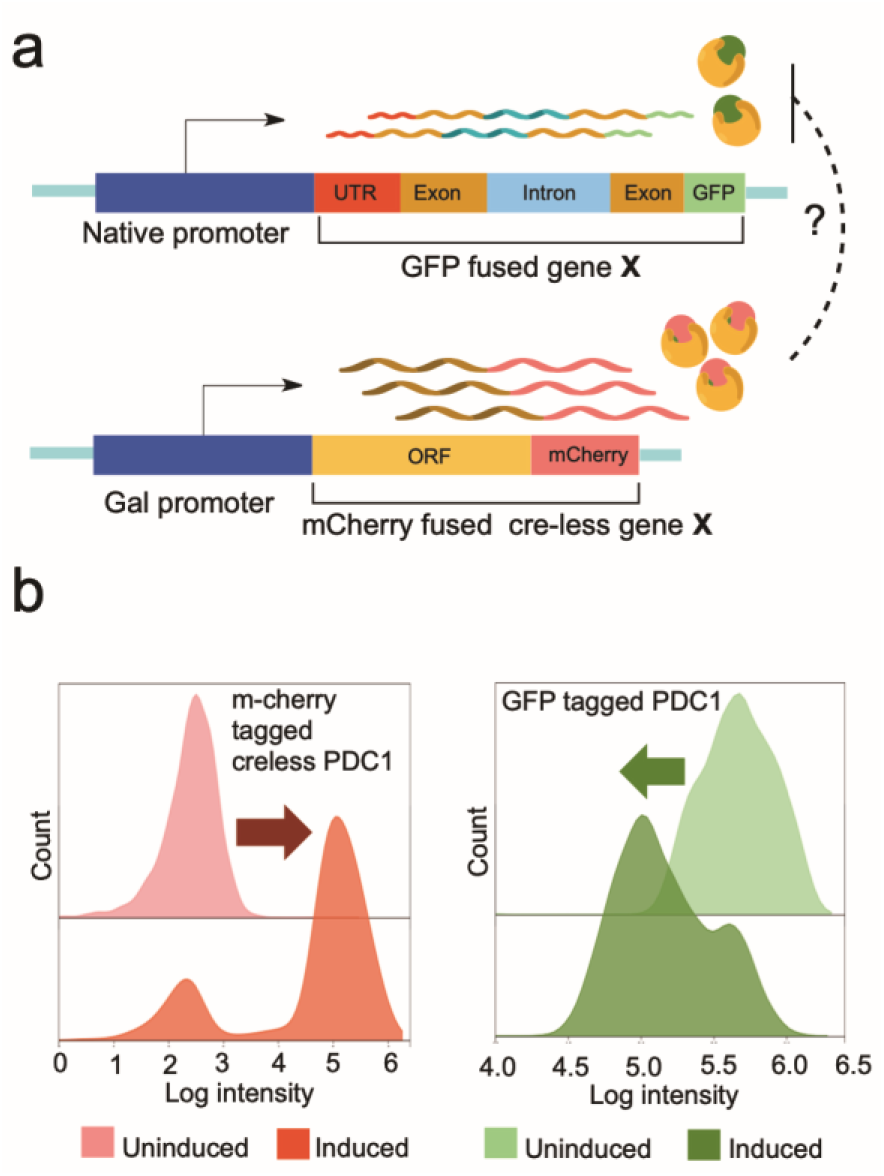
Screening for protein autoregulation. (**a**) One endogenous gene is fused to GFP and a cre-less (cis-regulatory element-less) version of the same gene is fused to mCherry. The cre-less version lacks the native 5’ and 3’ UTRs and any introns and has shuffled codons in order to prevent the protein autoregulating the cre-less mRNA. After inducing the cre-less gene, possible autoregulation is detected by a decrease in GFP levels. (**b**) An example of autoregulation with the gene PDC1. Overexpression of the cre-less PDC1 (seen as an increase in mCherry fluorescence) leads to a decrease of the endogenous PDC1 (seen as a decrease in GFP fluorescence).

The pyruvate decarboxylase gene *PDC1* is transcriptionally autoregulated ^65^. To test our strategy, we synthesized a cre-less copy of *PDC1*, fused to mCherry and under control of the *GAL1* promoter. Figure 1b shows the change in the fluorescence signal of both mCherry and GFP after 10 hours of induction, when the GFP signal is reduced about 10-fold. Note that there is a subset of cells that do not induce mCherry fluorescence, which results in the shoulder seen on the GFP fluorescence signal. To simplify measurements of the change in expression, in all further examples we use the measurement of GFP after induction only from cells that show induction of the mCherry signal.

### Screen of ribosomal proteins

We are primarily interested in identifying cases of post-transcriptional autoregulation, examples of which are common among ribosomal genes in bacteria ^27,29,30,50^. Among the known examples of yeast ribosomal genes that are post-transcriptionally autoregulated, repressing proper splicing of the pre-mRNA is often the mechanism ^56–61^. Many yeast ribosomal proteins have paralogs which are identical, or nearly so, to each other and for those cases it is sufficient to make a cre-less version from only one of the paralogs and to test the effects on expression of both wild-type (with GFP-fusion) paralogs. GFP expression of 60 large subunit ribosomal protein genes, including 25 paralogous pairs, both with and without induction of the cre-less gene, are shown in Figure 2a. Figure 2b shows the same for 35 small subunit ribosomal protein genes, including 16 paralogous pairs, and the fluorescence signal from control cells lacking a GFP-fusion gene. Shown are the median of the log (fluorescence) values of two or more measurements with the standard deviations also shown, which are generally quite small. Figure 2c shows both the induced and uninduced measurements in the same plot for all of the genes. Two genes, *RPL1B* and *RPS22B*, have greater than one log (>10-fold) decrease in expression after induction (marked with ** and above the 1-log dotted line), and another 19 genes have greater than ½ log change in expression (marked with *). One pair of paralogs, *RPL26A* and *RPL26B*, are both decreased by greater than ½ log, and a few others have similar changes of both paralogs, such as *RPL24A* and *RPL24B*, but only one exceeds the ½ log threshold. Cases where both paralogs are decreased may represent examples of increased rates of protein degradation of the overexpressed proteins ^52^. But in most cases only one paralog has a significant reduction in expression and is therefore likely to indicate direct repression of one gene’s expression. Of the known examples described above, we see a large reduction in expression of *RPL22B*. We did not see a large reduction in expression for *RPS14B*, but it is expressed at very low levels, consistent with previous reports that the ratio of *RSP14A* to *RSP14B* is 10:1 ^56^. We also did not see a large reduction in expression of *RPS28B,* likely because its regulation requires the 3’ UTR which is disrupted in the GFP-fusion genes^62,63^. This indicates one limitation of our approach, that the GFP fusion to the C-terminus of the protein disrupts the normal 3’ UTR, and cis-regulatory elements residing in that region will likely be missed. *RPL30* and *RPS9A*/*B* are missing from our GFP collection and could not be tested. The two genes we observe with the largest effects, *RPS22B* and *RPL1B*, have not, to our knowledge, been previously shown to be autoregulated, nor have most of the regulated examples we observe.

**Figure 2:**
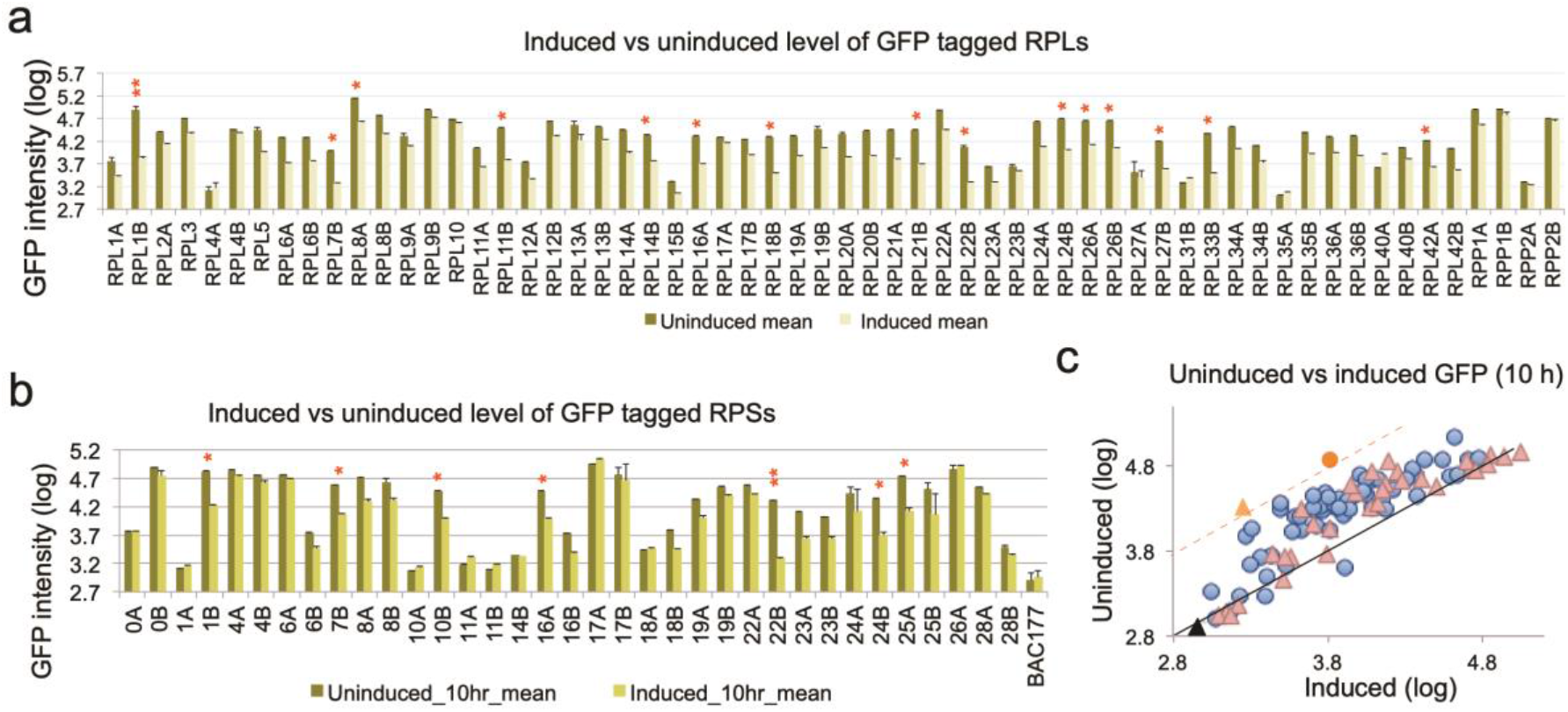
Screening of ribosomal protein autoregulation. (**a**) 60 large ribosomal subunit proteins were screened for autoregulation. Genes with more than a 10-fold reduction of their endogenous protein after induction of their cre-less protein are highlighted with two red stars (**). Genes with a 3-fold to 10-fold reduction are highlighted by one red star (*). (**b**) 35 small ribosomal subunit proteins were screened for autoregulation. BAC177 is a negative control with no GFP-tagged proteins. (**c**) Scatterplot showing the uninduced vs induced log10 GFP levels for large ribosomal subunit proteins (triangles) and small ribosomal subunit proteins (circles). *RPL1B* (red circle) and *RPS22B* (orange triangle) showed the highest levels of autoregulation. The black line passing through the center represents equal GFP levels for induced and uninduced, while the orange dotted line represents a 10-fold reduction in the induced compared to uninduced. The black triangle is BAC177 with no GFP-tagged proteins.

### Autoregulation of *RPS22B*

Upon induction of the cre-less version of *RPS22* (the two paralogs code for identical proteins), the expression of wild-type *RPS22B* (with GFP fusion) is reduced over 10-fold at 10 hours while *RPS22A* has only a modest reduction (Fig. 2b). The *RPS22A* gene has no introns whereas the *RPS22B* gene has two introns, including one in the 5’ UTR that contains a conserved, predicted secondary structure ^66^. That intron contains seven AUG codons and is a substrate for RNase III-mediated cleavage if not spliced ^67^, both of which suggest that splicing is required for translation of the mRNA. In fact, deletion of the 5’ UTR intron increases expression of the *RPS22B* gene several-fold ^68^. The simplest hypothesis is that the 5’ UTR intron of the RPS22B gene is the cis-regulatory site required for autoregulation, probably via inhibition of splicing. To test that we integrated the gene for GFP expressed from the constitutive *TEF2* promoter, whose activity is unaffected by galactose induction ^69^, into the yeast chromosome in place of a putative gene of unknown function (chromosome II: YBR032W). Two different 5’ UTRs were placed upstream of GFP, the complete *RPS22B* UTR and a “post-spliced” *RSP22B* UTR with the intron removed. Both strains were transformed with the plasmid containing the cre-less version of *RPS22B* (Figure 3a). GFP fluorescence was measured for each strain, with and without the plasmid, and with and without induction (Figure 3b and 3c). The strain with the spliced 5’ UTR showed nearly identical expression with or without the plasmid and with or without induction (Figure 3b and 3c, blue traces), indicating that translation of the spliced mRNA is not repressed. In the strain with the complete *RPS22B* 5’ UTR, GFP expression is nearly 3-fold lower in cells without the plasmid and with the plasmid but without induction. We expect this is due to repression by the endogenous Rps22 protein in the cells. When the cre-less plasmid is induced to overexpress Rps22, the expression of GFP is further reduced about 6-fold (figure 3b and 3c, green traces), similar to the repression of the wild-type *RPS22B*-GFP strain (Figure 2b). This indicates that, similar to several other autoregulated ribosomal protein genes, repression occurs by inhibiting splicing. Interestingly, the bacterial ortholog of Rps22 is S8 ^70^ and it is also involved in autoregulation ^49^. One anomaly is worth noting. The original cre-less version of Rps22 had unintentionally left the stop codon at the end of the gene, so that it was not fused with mCherry, but it showed the autoregulation. When that was corrected to make a cre-less version fused to mCherry, autoregulation was no longer observed, indicating interference with RNA binding by the mCherry fusion. It is possible the same thing happens with other of our mCherry fusion constructs, which is another reason we could be underestimating the true number of autoregulated ribosomal protein genes.

**Figure 3:**
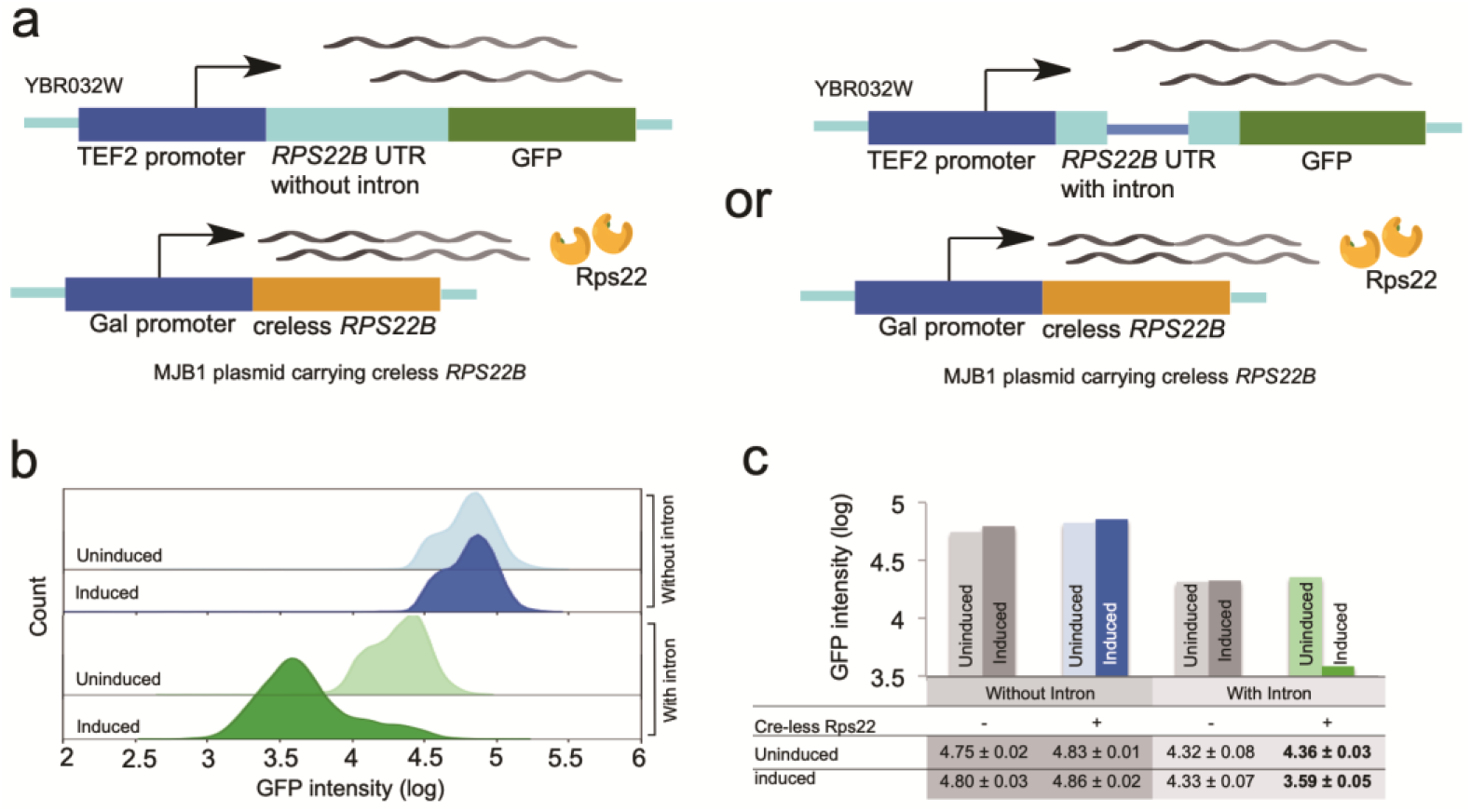
Regulation of RPS22B. (**a**) Two reporter constructs were made to test the sequence requirements for autoregulation by overexpressed cre-less Rps22. One construct had the wild-type *RPS22B* 5’ UTR (containing a 557 bp intron) placed upstream of GFP and driven by the *TEF2* promoter. A second construct was identical except that the intron was removed. Both constructs were separately integrated into the YBR032W locus and transformed with the plasmid for expressing cre-less Rps2. (**b**) Histograms of log GFP intensity from flow cytometry of cells containing either the UTR-with-intron reporter (green) or the UTR-without-intron reporter (blue). Cells were measured as either uninduced (lighter-shaded colors) or after induction of cre-less Rps22 (darker-shaded colors). Each histogram curve represents cells picked from a single colony after transformation of the cre-less plasmid. Multiple colonies for each reporter construct were picked and measured this way, and these replicates were used to generate the bar plot in 3c. (**c**) Bar plot comparing the log GFP levels of the two reporter constructs in cells with or without the plasmid for expression of cre-less Rps22 and with or without galactose induction. Only the UTR-with-intron construct (green) shows autoregulation after induction of cre-less Rps22.

### Autoregulation of *RPL1B*

The expression of the *RPL1B* gene is reduced over 10-fold after 10 hours of induction of the cre-less version (Figure 2a). The paralog *RPL1A* showed only about a 2-fold reduction in expression (Figure 2a). *RPL1B* has no introns and we surmised that the 5’ UTR may be the regulatory region. The gene has a short 5’ UTR of 64 bases that is highly conserved within the *senso stricto* yeast species ^71^ (Figure 4a). We place the full 5’ UTR before the chromosomal GFP gene (as we did for the *RPS22B* UTR described above) and measured about a 10-fold reduction in expression after induction of the cre-less gene (Supplementary figure S2, green graphs). This indicates that the 5’ UTR is sufficient to confer feedback regulation by Rpl1. We also tested different versions of the gene on the plasmid fused to mCherry. Besides the cre-less version (*RPL1B-cl*), we tested three alternative constructs (Supplemental figure S3): wild-type for both the 5’UTR and coding region (*RPL1B-wt*); wild-type for the 5’UTR with a shuffled coding region (*RPL1B-cd*); and mutant 5’UTR but wild-type coding region (*RPL1B-mt*). After induction the two constructs containing the wild-type 5’UTR (*RPL1B-wt* and *RPL1B-cd*) produced about 2-fold less mCherry fusion protein than those with the mutant 5’ UTR (*RPL1B-mt* and *RPL1B-cl*), consistent with the regulatory element being within the 5’ UTR (Supplementary figure S4). Although the effect is smaller, the constructs with the wild-type 5’ UTR also reduce the repression of GFP-tagged *RPL1B*, presumably as a result of elevated level of Rpl1-mCherry protein. The wild-type and cre-less versions of *RPL1B*-mCherry both reduce *RPL1A* expression equivalently, about 3-fold after 10 and 20 hours of induction (Supplementary figure S5). To further test the evidence for whether Rpl1 directly interacts with the 5’ UTR sequence of its own message we designed an EMSA experiment with purified Rpl1 (see Methods) and a synthetic RNA, containing the UTR sequence. The presence of a distinct slower migrating band in lanes 2-3 (Supplementary figure S6) indicates direct binding between Rpl1 and the 5’-UTR of its own message.

**Figure 4:**
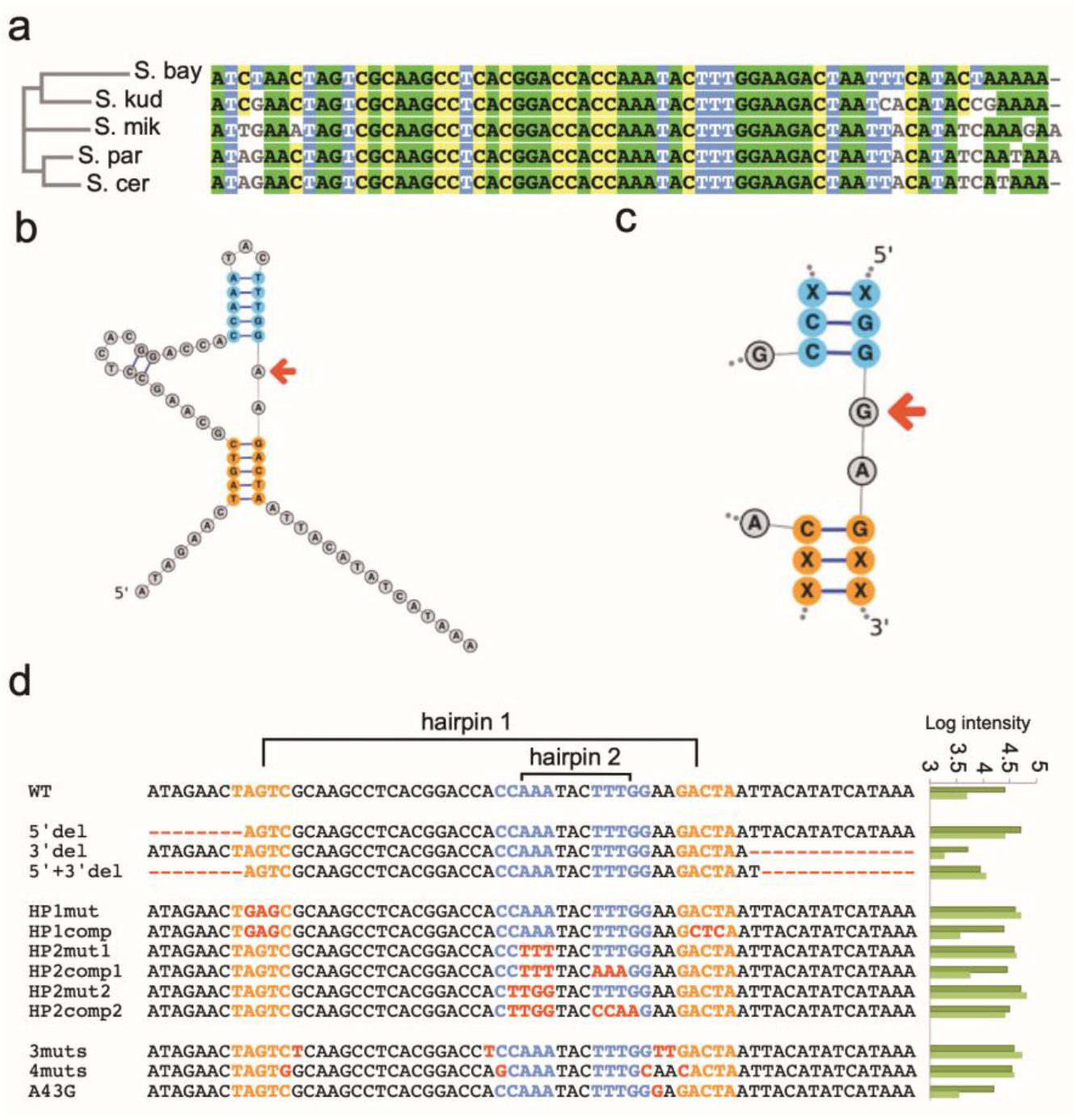
*RPL1B* autoregulation. (a) Alignment of the 5’ UTR sequence of *RPL1B* across different species of saccharomyces shows a high level of conservation. (b) The predicted structure of the *RPL1B* 5’ UTR with hairpin stem 1 (HP1) in orange and hairpin stem 2 (HP2) in blue. The red arrow indicates A at position 43 which differs from the G nucleotide seen in the consensus motif bound by Rpl1 in other species. (c) The common sequence and structure motif bound by Rpl1 in other species. The red arrow indicates the G nucleotide that is changed to A in *RPL1B*’s 5’ UTR sequence. (d) Variant sequences of the *RPL1B* 5’ UTR that were used to probe the sequence and structure requirements for autoregulation by Rpl1. Variants included deletions from the 5’ and 3’ ends, mutations to disrupt the hairpin stem structures as well as compensating mutations to restore them, and mutations affecting the core GGAAG of the motif. The log GFP expression for uninduced (dark green) and induced (light green) cells are shown to the right for each sequence.

Having verified that there is a post-transcriptional cis-regulatory element in the 5’ UTR of *RPL1B* that responds to the Rpl1 concentration, we sought to identify the sequence features required for binding by Rpl1. The RNAStructure web server ^72^ (see Methods) predicts the same minimum free energy secondary structure and maximum expectation secondary structure for the 64 base long 5’ UTR (Figure 4b). The structure has two primary stems, with hairpin stem 1 (HP1) shown in orange and hairpin stem 2 (HP2) shown in blue. Remarkably, every known binding site for Rpl1 and its orthologs in other species contains a common sequence and structure motif, shown in Figure 4c. This includes the binding sites for *E. coli* ribosomal protein L1 on both the L11 mRNA and the 23SrRNA ^73^, and ribosomal proteins from several other bacteria and archaea on mRNA and rRNA binding sites ^49,74^. It also includes the binding sites on stems H77 and H78 of 28SrRNA of both human and yeast ^75,76^. The *RPL1B* UTR structure contains the same sequence and structure motif except for a single G to A change shown by the arrows in Figures 4b,c.

To test if those sequence and structure features are required for regulation by Rpl1, we made several variants of the *RPL1B* 5’ UTR and placed them upstream of GFP driven by the *TEF2* promoter. Figure 4d shows the wild-type 5’UTR with the Hairpin1 and Hairpin2 stems marked. Below that are 12 different variants and the logarithm of the GFP expressions from cells both uninduced (-Gal) and induced (+Gal) for the expression of the cre-less Rpl1 protein. The log expression values and the differences between induced and uninduced are also shown in the Table 1. The wild-type UTR is repressed by 0.70 (log reduction in GFP), verifying that the 5’ UTR is the regulatory region. Removing the bases at either the 5’ or 3’ sides of the structured region reduces but does not eliminate repression, but removing both together does eliminate repression, perhaps by altering the structure of the mRNA. To test the importance of the hairpin structures we modified the 5’ half of each stem to eliminate the structure, and then compensated by modifying the 3’ half to make a complementary sequence and recover the secondary structure. HP1-mut modifies the 5’ half of the HP1 sequence to eliminate the structure and it completely abolishes repression. HP1-cmp restores the predicted secondary structure and regains regulatory activity, in fact to a slightly higher level than the wild-type sequence.

**Table 1.**
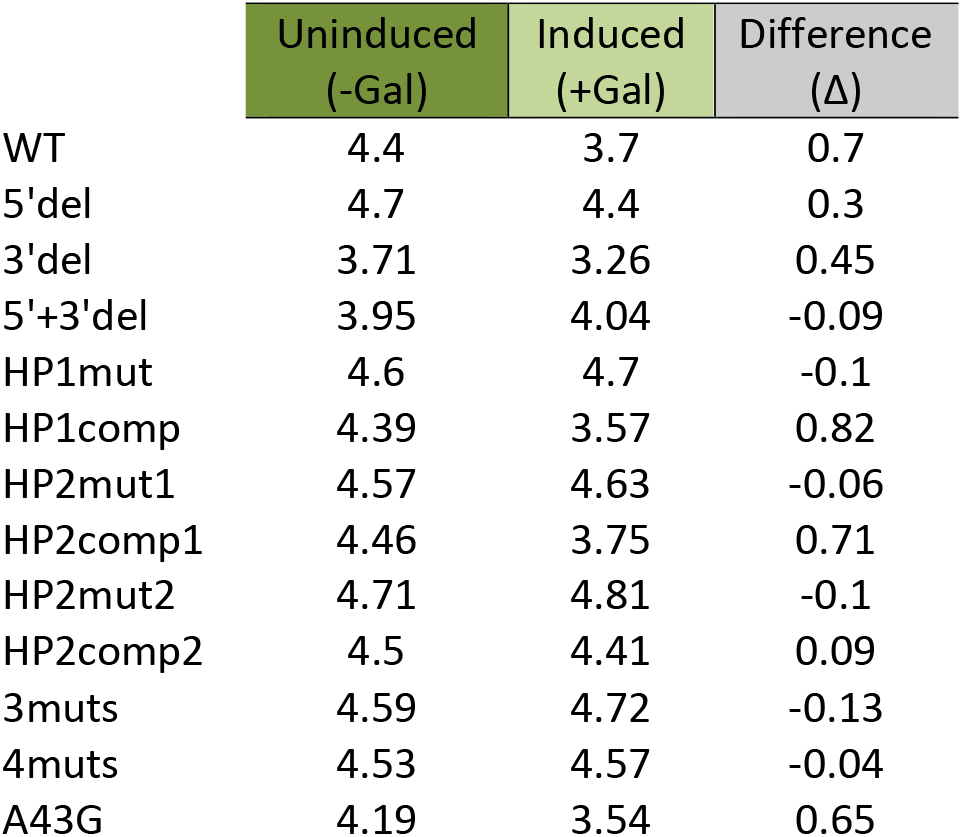
GFP expression level (log) with or without induction in the presence of *RPL1B* UTR variants.

We mutated HP2 in two different ways, first altering only the upper three bases in the 5’ half (HP2-mut1) and then by altering the upper four bases in the 5’ half (HP2-mut2). Both eliminate repression, demonstrating the importance of HP2 for the regulatory site. When the upper three bases are compensated (HP2-cmp1), repression is restored. However, when the upper four bases of HP2 are compensated (HP2-cmp2), there is still no regulation. This highlights the importance of the first G in the conserved sequence GGGAG shown in Figure 4c. To further test the importance of the bases in the conserved motif we altered the GGAAG to GGTTG (3muts, which also contains an additional mutation to maintain the wild-type structure), and the repression is eliminated. In another variant we changed the GGAAG to GCAAC, along with two additional changes to maintain the secondary structure (4muts), and again the repression is eliminated. Finally, we tested the importance of the base that differs between the *Rpl1B* 5’ UTR (the A with the arrow in Figure 4b) and the conserved G in the binding site motif (the arrow in Figure 4c). The A to G variant (A43G) is still repressed, indicating that either base is acceptable at that position for regulatory activity. However, the expression of GFP in the non-induced state is lower for the mutant A43G than for the wild-type sequence, perhaps because it provides a higher affinity binding site for the intrinsic Rpl1 protein in the cell.

## Conclusions

Many recent studies use high-throughput methods to identify protein-mRNA interactions, some of which may be regulatory interactions. However, most of those approaches will only identify common events, where a protein regulates many genes. Discoordination of protein and mRNA levels adds to the complexity of these studies and any inferences that can be drawn. Additionally, cases of autoregulation, such as in bacteria, would be missed because the proteins have only a single target and are not, primarily, regulatory proteins. Non-coding segments of RNA can evolve rapidly and acquire various roles including regulation of gene expression. They can be highly sensitive and autonomous sensors of the cellular environment and determine their own fate, such as to be translated or not. Riboswitches are an especially compelling example of such autonomous regulation because no protein is involved in the feedback response. Sensing protein concentrations can also be accomplished by RNAs and is likely to be much more wide spread than is currently known, but because of the single target limitation to detection, directed searches are needed to identify such cases. We have used the GFP-fusion collection in yeast as a means of rapidly screening for examples of feedback regulation. We observed that more than 20% of ribosomal genes appear to be autoregulated, and that is likely an underestimate of the true number due to some limitations of our approach. We find that *RPS22B* regulates its own splicing, which is required for translation and is similar to a few other known autoregulatory examples in yeast. *RPL1B*, on the other hand, appears to regulate its own translation via binding to a sequence and structure motif in the 5’ UTR that is remarkably conserved in bacteria, archea and eukaryotes from yeast to mammals. We know that motif is necessary but it doesn’t appear to be sufficient as changes outside of the conserved region can also affect regulation. More work is required to identify the complete mechanism, and to uncover the regulatory domains of the remaining examples we identified. The GFP-fusion collection in yeast provides an outstanding resource for identifying feedback regulation of all types, but the development of more flexible approaches will be necessary to do similar searches in more complex genomes.

## Materials and Methods

### GAL1 vector and yeast strains

The synthetic genes were cloned into a custom plasmid, MBJ1-mod5 (Supplementary figure S7). The vector backbone was pMW102-empty-MORF. The main features of this vector are a β-lactamase gene, replication origin for selection in E. coli, the URA3 gene for selection in ura-yeast strains and GAL1 promoter for induction with galactose and the mCherry coding region that can be fused to the cre-less gene. The His5 terminator site with 5’-NheI and 3’-XhoI restriction sites was synthesized by G-blocks from IDT. The mCherry sequence was cloned out from pMVS124-pACT1 (a generous gift from Max Staller) with a 3’-NheI and 5’-AvrII restriction sites by PCR using. pMW102-empty-MORF was linearized using primers to incorporate PacI and AvrII restriction sites downstream of the GAL1 promoter. One microgram of purified mCherry, His5 terminator and linearized vector backbone were digested with NheI, AvrII, XhoI and PacI for 15 min and gel purified. The digested DNAs were ligated with T4 DNA ligase for 15 min at room temperature and transformed into DH5α cells. The sequences of the selected clones were verified by Sanger sequencing.

The cre-less genes were synthesized and integrated directly into the MBJ1-mod5 vector by Twist Biosciences (San Francisco, CA). The genes were designed to be lacking any cis-regulatory elements (cre-less) by eliminating any introns, replacing the 5’ UTR with an alternative sequence, and shuffling the synonymous codons of the gene using the program CodonShuffle ^64^. The sequences of the shuffled coding regions are provided in the supplementary file2.

The parental GFP-tagged yeast strains (S288C) were taken from the Yeast GFP library ^48^ (a gift from Heather True-Kro). GFP-tagged strains were transformed with the plasmid containing the corresponding cre-less gene driven by the GAL1 promoter. Two independent clones were assayed for each sample.

### Yeast transformation with cre-less plasmid

Yeast strains from the GFP collection (*MATα SUC2 gal2 mal2 mel flo1 flo8-1 hap1 ho bio1 bio6*) were grown in YPD medium (1% yeast extract, 1% bacto-peptone and 2% glucose) at 30 ℃, overnight in 96 well plate format or in individual culture tubes. The cells were inoculated in 1 ml fresh YPD (10% v/v) and grown to OD_600_ = 1. The cells were collected by centrifugation at 2000 Xg for 2 min and then mixed with 0.6-1 μg of the cre-less plasmid in buffer containing 100 mM LiOAc, 50 % PEG (MW 3350). The resulting mixture was incubated for 5 min and subjected to heat shock at 40 ℃ for 20 min. The cells were mixed with 200 μl of fresh YPD, incubated at 30 ℃ with shaking and plated on selective medium (SD-URA). Several colonies from each plate were collected after 2-3 days for galactose induction.

### Galactose induction and flow cytometry assay

Liquid cultures were inoculated and grown overnight in 400ul SD-URA with 2% raffinose in 96 deep well plates at 30 ℃. Overnight cultures were diluted into both SD-URA with 2% raffinose (uninduced) and SD-URA with 2% raffinose and 0.2% galactose (induced). Cells were grown at 30C for 10 hours.

200 μL cultures were transferred to 96-well plates and assayed on a CytoFLEX (Beckman Coulter). The live cells were gated and 10,000 events were acquired.

### *RPL1B* Variants

Three additional *RPL1B* variants were synthesized (Supplementary figure S3) beside the cre-less version of the gene (*RPL1B-cl*). The *RPL1B-wt* retained the wild type sequence of native *RPL1B* with the 5’ UTR. The *RPL1B-cd* retained the 5’ UTR of the wild type sequence but the internal mRNA sequence was shuffled and was identical to the cre-less sequence. The *RPL1B-mt* was identical to the wild type coding sequence but lacked the 5’ UTR. All of these variations of *RPL1B* sequences were fused with mCherry sequence and cloned into MJB1 plasmid as described above.

### Reporter gene assay

To construct yeast strains expressing GFP with either the *RPS22B* UTR or *RPL1B* UTR variants a new plasmid (BAC690-TEF) was designed. BAC690-TEF (Supplementary figure S8) was designed based on the BAC690_Euroscarf vector, which has an eGFP ORF 5’ to an ADH1 terminator ^77^. The TEF2 promoter sequenced was amplified using PCR from purified yeast saccharomyces cerevisiae (S288C) genomic DNA. The TEF2 promoter was cloned into the BAC690_Euroscarf vector 5’ of the eGFP sequence. UTR variants were cloned upstream of eGFP using NEB’s HiFi DNA Assembly. A PCR amplicon was made containing the TEF2 promoter, UTR variant, eGFP, and ADH1 terminator as well as a kanamycin phosphotransferase cassette for integration in the yeast genome. The integration location was a dubious ORF (YBR032W).

Transformed cells were grown on YPD plates containing 200ug/ml G418 to select for the integrated kanamycin cassette. Correct integrations were verified by colony PCR of the 5’ and 3’ junctions. Integrated strains were then transformed with their corresponding cre-less plasmid. Galactose induction and flow cytometry measurements were done as described above.

### RNA Structure prediction

We used the Matthews lab <u>RNAstructure</u> prediction web server ^72^ (https://rna.urmc.rochester.edu/RNAstructureWeb/Servers/Predict1/Predict1.html) version 6.0.1 with default parameters. It produces both a minimum free energy predicted secondary structure and a maximum expectation prediction secondary structure, which may be different. The structures shown in Figures 4b,c were created using the StructureEdit program from the same package.

## Acknowledgements

Funding for this work came from NIH grants R01HG000249 and R21GM126307. FB was supported by R25HG006687. We thank Dr. Max Staller for the gift of a plasmid used in this work and Dr. Heather True-Krob for the gift of the yeast GFP-fusion collection.

