## supplemental file 1 for "Autoregulation of Many Yeast Ribosomal Proteins Discovered by Efficient Search for Feedback Regulation"

### Supplementary Information
### For

**Autoregulation of Many Yeast Ribosomal Proteins Discovered by Efficient Search for Feedback Regulation**

Basab Roy[1], David Granas[1], Fredrick Bragg, Jonathan Cher, Michael White, Gary D Stormo

Department of Genetics and Center for Genome Sciences and Systems Biology

Washington University School of Medicine, St Louis, MO 63110

[1] co-first authors

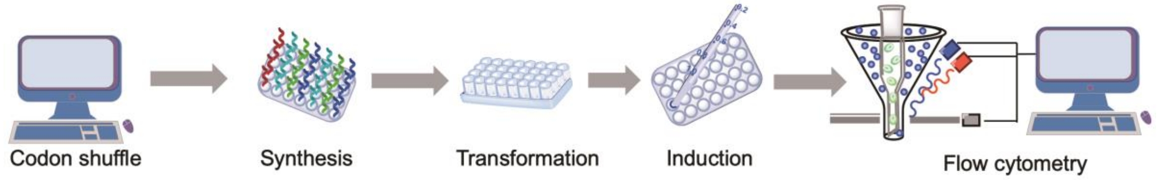


S1: Scheme for testing autoregulation. The cre-less genes were synthesized without cis regulatory elements, such as introns and UTRs, and their codons were shuffled to eliminate the chance of autoregulation by the cre-less protein. The cre-less genes were cloned into the MJB1 vector driven by the GAL1 promoter. Each cre-less gene plasmid was transformed into a yeast strain that had a GFP-tagged version of the corresponding gene at its native locus. Transformations were done either individually or in 96-well plate format. The cre-less protein was induced by adding galactose. The level of cre-less protein and the native protein were monitored by mCherry and GFP fluorescence, respectively.

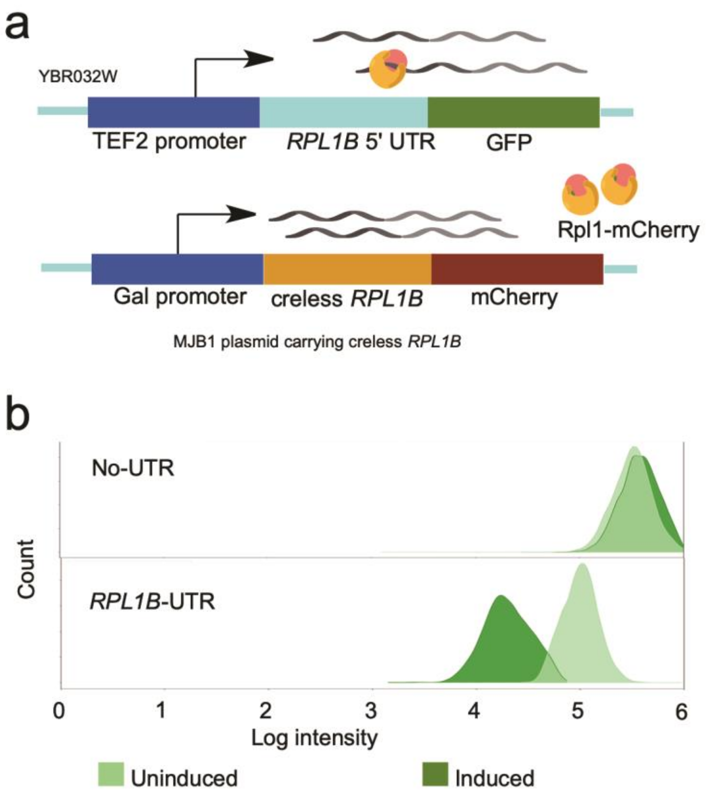


S2: Reporter gene assay with *RPL1B* UTR. (**a**) A reporter construct was made by placing the *RPL1B* 5’ UTR upstream of GFP and having it driven by the *TEF2* promoter. A negative control construct was also made without the *RPL1B* 5’ UTR. Each construct was integrated in the genome (Chromosome II, YBR032W) and the cells were transformed with the cre-less *RPL1B* plasmid. (**b**) The cre-less *RPL1B* was induced with galactose and GFP fluorescence was measured to detect autoregulation. A decrease in GFP after induction only occurred with the construct containing the *RPL1B* 5’ UTR showing it is sufficient to allow autoregulation by Rpl1.

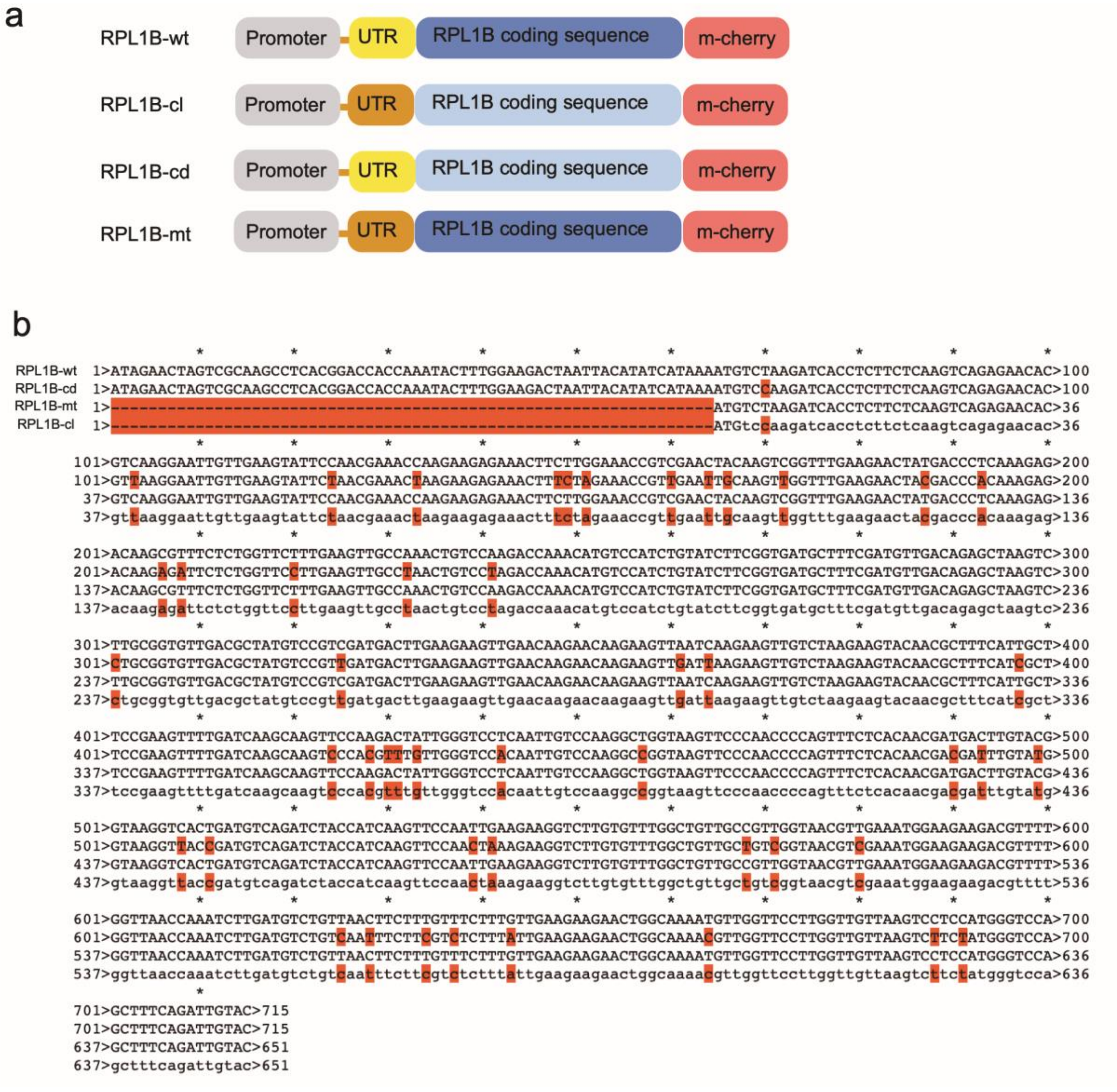


S3: Design and sequence alignment of *RPL1B* variants used to compare the contribution of regulatory elements in protein autoregulation. Beside the cre-less version (RPL1B-cl) three other alternative structures were synthesized: wild-type for both the 5'UTR and coding region (RPL1B-wt); wild-type for the 5'UTR with a shuffled coding region (RPL1B-cd); and mutant 5'UTR but wild-type coding region (RPL1B-mt). All of the three variants were tagged with mCherry and cloned in the MJB1 vector for expression by galactose induction. (**a**) Cartoon of the *RPL1B* gene constructs with different elements, where differences in sequences within each region are indicated by altered colors. (**b**) Sequence alignments of the four *RPL1B* constructs with mismatches highlighted in red.

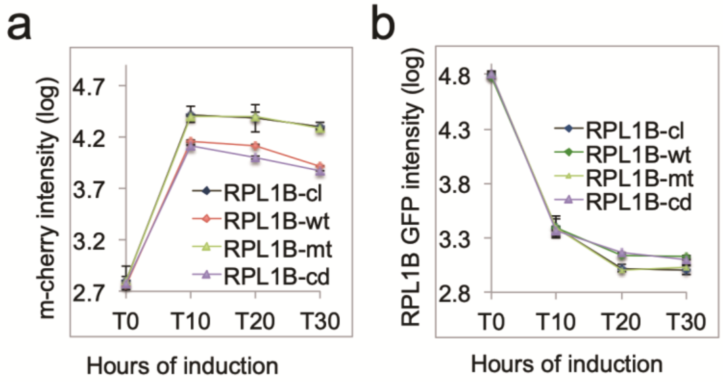


S4: The mCherry and GFP expression levels of RPL1B-cl, RPL1B-wt, RPL1B-cd and RPL1B-mt after 10, 20 and 30 hours of galactose induction. (**a**) The mCherry signal is indicative of RPL1B-cl, RPL1B-wt, RPL1B-cd and RPL1B-mt from the MJB1 plasmid. (**b**) The GFP signal indicates the expression level of the chromosomal copy of *RPL1B*

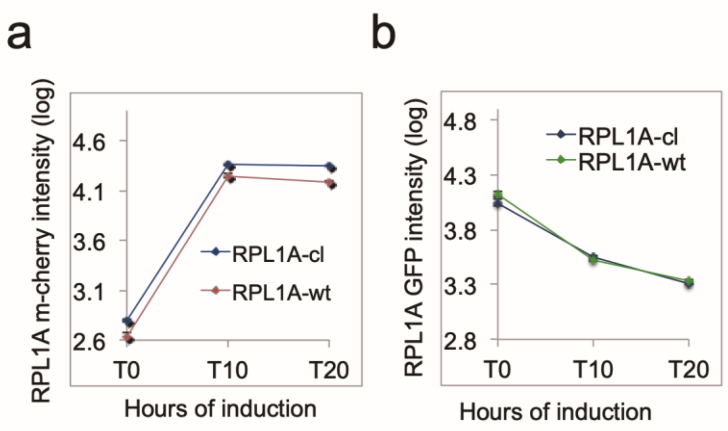


S5: The mCherry and GFP expression levels of RPL1A-cl and RPL1A-wt after 10 and 20 hours of galactose induction. (**a**) The mCherry signal is indicative of RPL1A-cl and RPL1A-wt from the MJB1 plasmid. (**b**) The GFP signal indicates the expression level of the chromosomal copy of *RPL1A*.

```
RPL1B-UTR   ATAGAACTAGTCGCAAGCCTCACGGACCACCAAATACTTTGGAAGACTAATTACATATCATAAAATGTCT

Control-UTR AGTAACAAAAAATTAAAGTTAATTAAGGAGATTAAACTATATATCAACAAAAAATTGTTAATATACCTCT
```

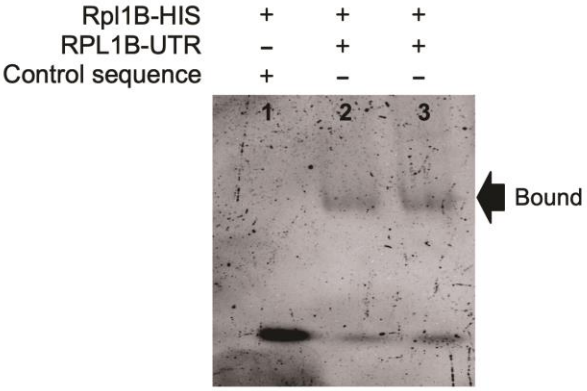


S6: EMSA with purified his-tagged Rpl1 (Rpl1B-HIS) and either the control sequence (lane 1), synthetic *RPL1B*-UTR RNA (lane 2) or *in-vitro* transcribed *RPL1B*-UTR (lane 3).

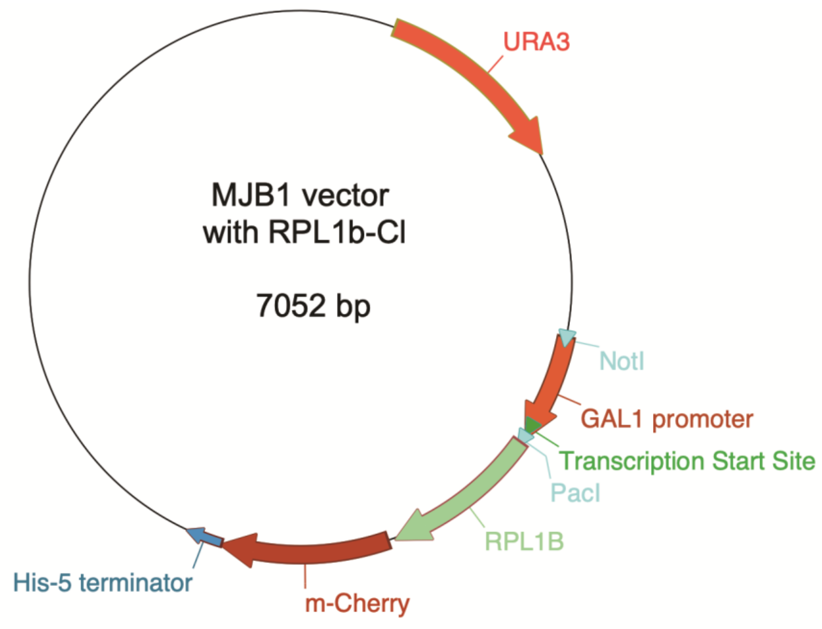


S7: The MJB1 vector with the expression cassette including GAL1 promoter, mCherry fused cre-less *RPL1B* (RPL1B-cl) and His-5 terminator.

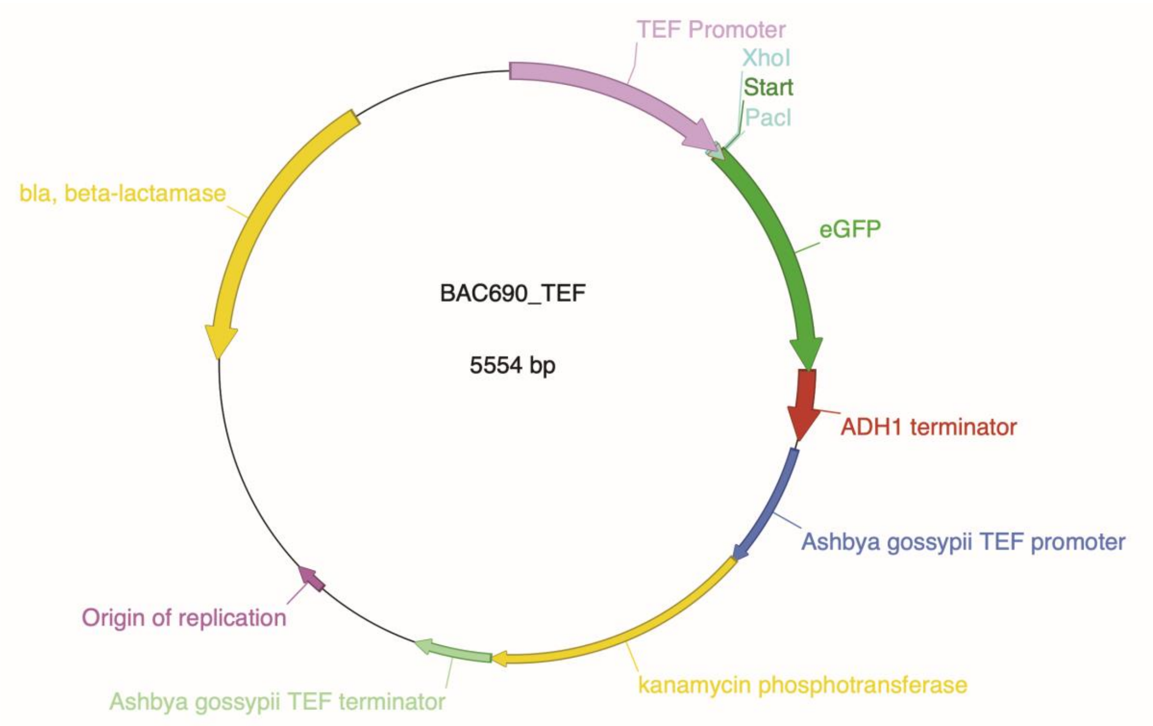


S8: The BAC690_TEF vector map showing the TEF2 promoter, cloning location (between XhoI and PacI sites) for placing UTRs from either *RPS22B* or *RPL1B* upstream of GFP and the ADH1 terminator. The whole cassette and the kanamycin phosphotransferase was amplified with flanking regions homologous to the site of insertion in the yeast genome (YBRO32W).

#### Supplementary Methods

##### Protein purification

The cDNA sequence of Rpl1B was codon optimized for expression in E. coli and cloned into SBP-PRDM vector (Ref A). The ORF was cloned downstream of the T7 promoter and was flanked by EcoRI and NotI restriction sites. An 18-nucleotide (CACCATCACCATCACCAT) sequence was attached to the 3′-end of the ORF for encoding an additional six amino-acid (HHHHHH) long peptide (His-tag) for purification. Expression of Rpl1B was achieved in E. Coli BL21 (DE3) cells. 250 ml E. Coli culture was grown in LB media until OD600 = 0.7 and induced by IPTG to 1mM final concentration for 18 hr. After lysing the cells by sonication, proteins were separated from the cellular debris by centrifugation at 15,000 rpm. The supernatant was filtered and loaded directly on a column prepacked with HisPur Ni-NTA resin (Thermo Scientific). After washing the column with buffer containing 50 mM Tris.HCl, pH 8.5 and 20 mM imidazole the protein was eluted in buffer containing 50 mM Tris.HCl, pH 8.5, 350 mM KCl, 20 mM $MgCl_2$ and 200 mM imidazole. The protein was stored in a final binding buffer with 50 mM Tris.HCl, pH 8.5, 350 mM KCl, 20 mM $MgCl_{2,}$ 0.1% NP-40 and 2% Glycerol. Buffer exchange was performed by spinning the sample at 400 rpm using Amicon ultra-4 centrifugal filter unit with a 10 kDa molecular weight cutoff. The protein was visualized by running a 4-15% Tris-glycine PAGE gel with 0.1% SDS at 200 V for 1h at room temperature. The gels were stained by Instant*Blue* (Sigma) and visualized by a Bio-Rad ChemiDoc imaging system.

##### RNA substrate production

Wild type *RPL1B*-UTR was cloned in a pUC19 vector under a T7 promoter. A negative control sequence of the same length was also cloned into the vector (Figure 4E, Supplementary figure S6). DNAs were amplified by PCR following the Phusion High-Fidelity PCR (NEB, catalog # E0553L) protocol. One microgram of the PCR product (linearized duplex DNA with 5 'T7 promoter and UTR or the control DNA) was used in a 50 μl *in vitro* transcription reaction following manufacturer's protocol (NEB). The mixture of DNA and enzyme was incubated at 37 ℃ for 16 hr. The RNA mixture was purified by Monarch RNA cleanup kit. The RNA concentration was measured by NanoDrop and adjusted to a 400 nM final concentration for EMSA.

##### Electrophoretic mobility shift assay

The protein−DNA binding reactions were done in buffer containing 50 mM Tris.HCl at pH 8.5, 350 mM KCl, 20 mM $MgCl_{2,}$ 0.1% NP-40, and supplemented with 10% glycerol. Either control RNA or *RPL1B*- UTR RNAs at a final concentration of 400 nM were incubated with 0.1 to 1 μg of purified Rpl1B for 30 min in 10 μL reaction volumes at 4 ℃. The reaction mixtures were run on a 7.5% Tris-glycine PAGE gel at 80 V for 2 hr in the cold room. The gels were stained with SYBR Green I stain for 20 min. The RNA in the bound (slow migrating) and unbound (fast migrating) bands were visualized by a BioRad imager with a 520 nM bandpass filter.
