## supplemental file 2 for "Autoregulation of Many Yeast Ribosomal Proteins Discovered by Efficient Search for Feedback Regulation"

Sequences of Codon-shuffled yeast genes

>RPS8A, YBL072C, SGDID:S000000168

ATGGGAATAAGCAGAGACTCTAGACACAAGCGTTCCGCTACTGGTGCTAAGAGAGCCCAATTCCGTAAGAAGAGA
AAGTTTGAATTGGGTAGACAACCAGCTAACACTAAGATTGGTGCCAAAAGAATCCACTCTGTTAGAACCCGTGGT
GGTAACAAGAAGTACAGAGCTTTAAGAATTGAAACTGGTAACTTCAGCTGGGCTTCTGAAGGCATCTCTAAGAAG
ACCAGAATCGCTGGTGTCGTCTACCACCCAAGCAATAACGAATTAGTTAGAACCAACACTCTAACCAAGGCCGCTA
TTGTTCAAATTGATGCTACCCCATTCCGTCAATGGTTCGAAGCCCATTACGGTCAAACCTTGGGTAAGAAGAAGAA
CGTTAAGGAAGAAGAAACTGTTGCTAAGTCTAAAAACGCTGAAAGAAAGTGGGCTGCTAGAGCTGCTTCCGCCAA
GATCGAATCCTCTGTTGAATCCCAATTCTCTGCCGGTAGATTGTACGCTTGTATTTCCTCCAGACCAGGTCAATCTG
GTAGATGTGATGGTTACATCTTGGAAGGTGAAGAACTAGCTTTCTACTTAAGAAGATTGACTGCTAAAAAGTAG

>RPS1A, YLR441C, SGDID:S000004433

ATGGCAGTTGGTAAAAACAAGAGATTGTCTAAAGGTAAGAAAGGTCAAAAAAAGAGAGTTGTCGATCCATTCACC
AGAAAGGAATGGTTCGATATTAAGGCTCCATCAACTTTCGAAAACAGAAATGTTGGTAAGACTTTGGTCAATAAGT
CCACCGGTTTAAAATCCGCTTCCGACGCTTTAAAGGGTAGAGTTGTTGAAGTCTGCTTGGCTGATTTGCAAGGTTC
TGAAGACCATTCCTTCAGAAAGATTAAGTTGAGAGTTGACGAAGTTCAAGGAAAGAACTTGTTGACCAATTTCCAC
GGAATGGACTTTACTACTGACAAGCTAAGATCTATGGTTAGAAAGTGGCAAACTTTGATCGAAGCCAACGTCACTG
TGAAGACTTCCGATGACTACGTCTTGAGAATCTTCGCTATCGCCTTTACCAGAAAGCAAGCTAACCAAGTTAAAAG
ACACTCTTACGCTCAAAGTTCCCATATCAGAGCTATTAGAAAGGTTATTTCTGAAATCTTGACCAAGGAAGTTCAG
GGTTCCACCTTGGCTCAATTGACTTCTAAGTTGATCCCAGAAGTCATCAACAAGGAAATTGAAAACGCCACCAAGG
ATATCTTTCCACTACAAAACATCCACGTTAGAAAGGTTAAGTTGTTGAAGCAACCAAAGTTCGACGTTGGTGCTTTA
ATGGCTTTACACGGTGAAGGTTCTGGTGAAGAAAAGGGTAAAAAGGTTACTGGTTTCAAGGACGAAGTTTTGGAA
ACTGTTTAA

>RPS6A, YPL090C, SGDID:S000006011

ATGAAACTTAATATATCCTACCCTGTCAACGGTTCTCAAAAGACTTTCGAAATCGACGATGAACACAGAATCAGAG
TTTTCTTCGATAAGAGAATTGGTCAAGAAGTTGATGGTGAAGCTGTTGGTGACGAATTCAAGGGTTACGTTTTCAA
GATTTCTGGTGGGAACGATAAGCAAGGTTTCCCAATGAAGCAAGGTGTCTTGTTGCCAACTCGTATTAAGTTGCTA
TTGACTAAGAACGTTTCTTGTTACAGACCACGTAGAGATGGTGAAAGAAAGAGAAAGTCTGTTAGAGGTGCCATC
GTTGGTCCAGACTTGGCTGTCTTGGCTTTGGTTATCGTTAAGAAGGGTGAACAAGAATTGGAAGGTTTGACCGAT
ACTACCGTCCCAAAGAGATTGGGTCCAAAGAGAGCTAACAACATCAGAAAGTTCTTCGGTTTGTCCAAGGAAGAC
GACGTCAGAGATTTCGTCATCCGTAGAGAAGTCACCAAGGGTGAAAAAACTTACACTAAGGCTCCAAAGATCCAA
AGATTGGTCACCCCACAAAGATTGCAAAGAAAGAGACACCAAAGAGCTTTGAAGGTCAGAAACGCTCAAGCTCAA
AGAGAGGCTGCTGCCGAGTACGCTCAATTGTTGGCTAAGAGATTGTCTGAAAGAAAGGCTGAAAAGGCTGAAAT
TCGTAAGAGAAGAGCCTCTTCTTTGAAGGCTTAA

>RPS9A, YPL081W, SGDID:S000006002

ATGCCCCGCGCTCCCCGTACATACTCTAAGACCTACTCTACACCAAAGAGACCTTACGAATCTTCTAGATTGGACGC
CGAATTGAAGTTGGCTGGTGAATTCGGTTTAAAGAATAAGAAGGAAATCTATAGAATTTCTTTCCAATTATCCAAG
ATTAGAAGAGCCGCAAGGGATTTGTTGACTAGAGACGAAAAGGATCCAAAGAGATTGTTCGAAGGTAACGCGTT
AATTAGACGTTTAGTCAGAGTCGGTGTTTTGTCCGAAGACAAGAAGAAGTTGGATTACGTTTTGGCTTTGAAGGTC

GAAGATTTCTTGGAGAGGAGACTACAAACTCAGGTCTATAAGTTAGGTTTGGCTAAGTCCGTTCACCACGCTAGA
GTTCTAATTACACAAAGACACATTGCCGTCGGTAAGCAAATTGTCAATATTCCCTCTTTCATGGTTAGGTTAGACTC
TGAAAAGCATATCGATTTTGCACCAACTTCTCCATTCGGTGGTGCTCGTCCAGGCAGAGTTGCTAGAAGAAACGCA
GCCCGTAAGGCCGAAGCTTCTGGTGAAGCTGCTGATGAAGCAGACGAAGCTGACGAAGAGTAA

>RPS0A, YGR214W, SGDID:S000003446

ATGAGTCTTCCGGCAACCTTCGATTTGACCCCAGAAGATGCTCAATTATTGTTGGCTGCCAACACCCACTTAGGTGC
CAGAAACGTCCAAGTTCACCAAGAACCATACGTTTTCAACGCTAGACCAGACGGTGTTCACGTTATCAACGTTGGT
AAGACTTGGGAAAAGTTAGTTTTGGCTGCAAGAATTATCGCCGCCATCCCAAACCCAGAAGACGTCGTTGCCATCT
CATCTAGAACTTTCGGTCAAAGGGCTGTTTTGAAGTTCGCTGCCCACACTGGTGCTACTCCAATCGCTGGTAGATTC
ACTCCTGGTTCTTTCACTAACTACATTACCAGATCTTTCAAGGAGCCAAGACTTGTTATCGTTACCGATCCAAGATCC
GATGCTCAAGCTATCAAGGAAGCTTCCTACGTCAACATTCCAGTTATCGCTCTTACCGACTTGGATTCCCCATCAGA
ATTTGTCGACGTCGCTATCCCTTGTAACAATAGAGGTAAGCACTCTATCGGTTTGATTTGGTATTTACTAGCCAGAG
AAGTCTTAAGATTGAGAGGTGCTTTGGTTGACCGTACTCAACCATGGTCTATCATGCCAGACCTATATTTCTACAGA
GATCCAGAGGAAGTTGAACAACAAGTTGCTGAAGAGGCTACCACCGAAGAAGCTGGTGAAGAAGAAGCTAAAGA
AGAAGTTACTGAAGAACAAGCCGAAGCTACCGAATGGGCTGAAGAAAATGCTGACAACGTTGAATGGTAA

>RPS25A, YGR027C, SGDID:S000003259

ATGCCACCTAAACAGCAGTTGTCTAAAGCTGCTAAGGCTGCCGCCGCTTTAGCTGGTGGTAAGAAGTCTAAGAAG
AAGTGGTCTAAGAAGTCCATGAAGGACAGAGCTCAACACGCCGTTATCTTGGACCAAGAAAAATACGACAGAATC
TTGAAGGAAGTCCCAACTTACAGATACGTCTCCGTCTCTGTTTTAGTTGACAGATTAAAGATTGGTGGTTCCCTTGC
TAGAATCGCTTTGAGACACTTAGAAAAGGAGGGTATTATCAAGCCAATCTCTAAGCACTCTAAACAAGCTATTTAC
ACTAGAGCTACCGCTTCCGAATAA

>RPS16A, YMR143W, SGDID:S000004751

ATGTCCGCCGTTCCTTCTGTTCAAACTTTCGGTAAGAAGAAATCTGCTACTGCTGTTGCCCATGTTAAAGCTGGTAA
GGGTTTGATCAAGGTCAACGGTTCCCCAATCACTTTGGTTGAACCAGAAATCTTGAGATTCAAGGTTTACGAACCA
TTATTGTTGGTCGGTTTAGACAAATTCTCCAACATCGATATTAGAGTTAGAGTTACTGGTGGTGGTCATGTTTCTCA
AGTTTACGCTATTCGTCAAGCTATTGCCAAGGGTTTAGTCGCTTACCATCAAAAATATGTTGATGAACAATCAAAAA
ATGAATTGAAGAAGGCCTTTACCAGTTACGATAGAACTTTATTGATCGCTGACTCCCGTAGACCAGAACCAAAGAA
ATTCGGTGGTAAGGGTGCTAGATCTAGATTCCAAAAGTCATACAGATAA

>RPS7B, YNL096C, SGDID:S000005040

ATGTCTTCCGTTCAGTCTAAAATTTTGAGCCAAGCTCCATCTGAATTGGAATTGCAAGTCGCTAAAACCTTCATCGA
CTTGGAATCTTCCTCTCCAGAATTGAAGGCAGATTTACGTCCACTACAAATCAAATCCATTCGTGAGATTGACGTTA
CAGGTGGTAAGAAAGCTCTAGTTTTGTTTGTCCCAGTTCCAGCTTTGAGCGCCTACCATAAGGTCCAAACTAAATTA
ACTAGAGAATTGGAAAAAAAGTTCCCAGACAGACACGTCATCTTCTTAGCAGAAAGAAGAATCCTTCCAAAGCCAT
CCAGAACCTCCAGACAAGTTCAAAAGAGACCTAGATCTAGAACCTTGACCGCTGTTCATGACAAGGTCTTGGAAG
ACATGGTCTTTCCAACCGAAATTGTTGGTAAGAGAGTTAGATATTTGGTCGGTGGCAACAAGATCCAAAAGGTCCT
ATTGGACTCCAAGGATGTTCAACAAATCGATTACAAGTTGGAAAGTTTCCAGGCTGTCTACAACAAGTTGACTGGT
AAACAAATTGTTTTCGAAATCCCATCTCAAACTAACTAA

>RPS4A, YJR145C, SGDID:S000003906

ATGGCCCGTGGTCCGAAAAAGCACTTGAAGAGATTGGCTGCCCCACACCACTGGTTGCTAGACAAGTTATCCGGT
TGTTACGCTCCAAGACCTTCTGCAGGTCCACATAAGTTACGTGAATCCTTGCCATTGATTGTTTTCTTGCGTAACCGT
TTAAAGTACGCCTTGAACGGTAGAGAAGTTAAGGCTATTTTGATGCAAAGACACGTTAAGGTCGATGGTAAGGTC
AGAACTGATACCACTTACCCAGCCGGTTTCATGGACGTGATCACCTTGGACGCTACTAATGAAAACTTCCGTCTAG
TCTACGATGTTAAGGGTAGATTTGCTGTTCACAGAATCACCGATGAAGAAGCTTCTTACAAGTTGGGCAAGGTCAA
GAAGGTCCAATTAGGTAAGAAGGGTGTCCCATATGTCGTCACTCACGATGGAAGAACCATCAGATACCCAGATCC
AAACATTAAGGTTAATGACACCGTTAAGATCGACTTAGCTTCTGGTAAGATCACTGACTTCATCAAGTTCGATGCT
GGTAAGTTGGTCTACGTTACTGGTGGTCGTAATTTGGGTAGAATTGGTACTATCGTCCACAAAGAAAGACACGAC
GGTGGTTTCGACTTGGTTCACATCAAGGATTCTTTGGATAACACTTTCGTCACTAGACTAAACAACGTTTTCGTTAT
CGGTGAACAAGGTAAACCATACATCTCCTTACCAAAGGGTAAAGGTATCAAGTTGTCTATCGCCGAAGAAAGAGA
CAGACGTAGAGCTCAACAAGGTTTGTAA

>RPS24A, YER074W, SGDID:S000000876

ATGTCCGATGCAGTCACGATCAGAACTCGTAAGGTCATCTCTAACCCATTGTTAGCTAGAAAGCAATTCGTCGTCG
ACGTTTTACACCCAAACCGTGCTAACGTTTCTAAGGATGAATTGCGTGAAAAGTTGGCTGAAGTCTACAAGGCCGA
AAAGGATGCTGTTTCCGTCTTTGGTTTCAGAACTCAATTCGGTGGTGGTAAGTCTGTTGGTTTCGGTTTGGTCTACA
ACTCCGTTGCCGAAGCTAAGAAGTTCGAACCAACTTACAGATTGGTCAGATACGGTTTGGCTGAAAAGGTTGAAA
AAGCTTCCAGACAACAAAGAAAGCAAAAGAAGAATAGAGACAAGAAGATCTTCGGTACCGGTAAGAGATTGGCC
AAGAAGGTTGCTAGACGTAACGCTGACTAA

>RPS19A, YOL121C, SGDID:S000005481

ATGCCGGGGGTCTCTGTAAGAGATGTCGCCGCTCAAGATTTCATTAACGCCTACGCTTCTTTCTTGCAACGTCAAG
GTAAGTTGGAAGTCCCAGGTTACGTTGACATCGTTAAGACTTCTAGCGGTAACGAAATGCCACCACAAGACGCTG
AAGGTTGGTTCTACAAGAGAGCTGCTTCTGTTGCTAGACACATTTACATGAGAAAACAAGTTGGTGTTGGTAAATT
GAACAAGTTGTACGGTGGTGCCAAATCCAGAGGTGTTAGACCATACAAGCACATCGACGCTTCTGGTTCTATCAAC
CGTAAGGTTCTACAAGCCTTAGAAAAGATTGGTATTGTTGAAATCTCTCCAAAGGGTGGTAGAAGAATTTCCGAAA
ATGGTCAAAGAGATTTGGACAGAATTGCTGCCCAAACCTTGGAAGAAGATGAATAA

>RPS23A, YGR118W, SGDID:S000003350

ATGGGCAAGGGCAAACCTAGAGGTTTGAACTCTGCTAGAAAGTTGAGAGTTCATAGAAGAAACAACCGTTGGGCT
GAAAACAATTACAAGAAGAGATTGTTGGGTACTGCTTTCAAATCTTCTCCATTTGGTGGTTCCTCTCACGCTAAGG
GTATCGTCTTGGAAAAGTTGGGTATCGAATCCAAGCAACCAAACTCTGCTATCAGAAAGTGTGTTCGTGTCCAACT
AATTAAGAACGGTAAAAAGGTTACTGCCTTCGTCCCAAACGATGGTTGTCTAAACTTCGTCGACGAAAACGATGAA
GTTTTGTTGGCCGGTTTCGGTAGAAAAGGTAAGGCAAAGGGTGATATCCCAGGTGTCAGATTCAAGGTTGTCAAG
GTCTCTGGTGTCTCATTGTTGGCCTTATGGAAAGAAAAGAAGGAAAAGCCAAGATCTTAA

>RPS18A, YDR450W, SGDID:S000002858

ATGTCCTTGGTCGTACAGGAACAAGGTTCTTTCCAACACATCTTGCGTTTATTGAATACCAACGTTGACGGTAACAT
CAAAATTGTCTACGCTTTGACTACCATCAAGGGTGTTGGTAGAAGATACTCCAACTTGGTTTGTAAAAAGGCTGAT
GTTGATTTGCATAAGCGTGCTGGTGAATTGACCCAAGAAGAATTGGAACGTATCGTTCAAATTATGCAAAACCCAA
CTCACTACAAAATCCCAGCTTGGTTCTTGAACAGACAAAACGATATTACTGATGGTAAGGACTACCACACCTTGGC
TAACAACGTTGAATCCAAGTTAAGAGACGACTTAGAACGTTTGAAGAAGATCAGAGCTCACCGTGGTATTAGACA
CTTCTGGGGTTTAAGAGTTAGAGGTCAACACACTAAGACTACTGGTAGACGTAGAGCTTAA

>RPS10A, YOR293W, SGDID:S000005819

ATGTTAATGCCCAAAGAGGACAGAAACAAGATTCACCAATACTTGTTCCAAGAAGGTGTTGTTGTTGCTAAGAAG
GACTTCAACCAAGCCAAGCACGAAGAAATTGATACTAAGAACTTGTATGTCATTAAGGCCTTGCAATCCTTGACCT
CAAAGGGTTACGTTAAGACCCAATTCTCTTGGCAATACTACTATTACACTTTAACTGAAGAAGGTGTCGAATACTTG
AGAGAATACTTGAACTTGCCAGAACACATCGTCCCAGGTACTTACATTCAAGAAAGAAACCCAACCCAAAGACCAC
AAAGAAGATACTAA

>RPS17A, YML024W, SGDID:S000004486

ATGGGCCGTGTCCGTACGAAGACTGTTAAGCGTGCCTCTAAAGCTTTGATCGAAAGATACTACCCAAAGTTAACCT
TGGACTTCCAAACTAACAAGAGATTGTGTGATGAAATCGCTACCATCCAATCTAAGAGACTTAGAAACAAGATTGC
TGGTTACACCACTCATTTGATGAAACGTATCCAAAAGGGTCCAGTTAGAGGTATCTCTTTCAAGTTGCAAGAAGAA
GAAAGAGAAAGAAAGGACCAATACGTTCCAGAAGTTTCTGCTTTGGACTTGTCTAGATCTAACGGTGTTTTGAACG
TTGACAACCAAACCTCTGATTTGGTTAAGTCCTTGGGTTTGAAGTTGCCATTGTCTGTCATTAACGTTTCTGCCCAAC
GTGACAGAAGATATAGAAAGCGTGTCTAA

>RPS26A, YGL189C, SGDID:S000003157

ATGCCCAAAAAACGTGCCTCCAACGGTAGAAACAAAAAGGGTAGAGGTCACGTCAAGCCTGTCAGATGTGTCAAC
TGTTCCAAGTCCATCCCAAAGGATAAGGCTATCAAGAGAATGGCTATTAGAAACATTGTTGAAGCCGCTGCTGTTA
GAGACTTGTCCGAAGCTTCTGTCTACCCTGAATACGCCTTACCAAAAACTTACAACAAGTTACACTACTGTGTTTCC
TGTGCTATTCACGCTAGAATTGTCAGAGTCAGATCTAGAGAAGATAGAAAGAACAGAGCTCCACCACAAAGACCT
AGATTCAACAGAGAAAACAAGGTCTCTCCAGCCGATGCTGCCAAGAAGGCTTTGTAA

>RPS22A, YJL190C, SGDID:S000003726

ATGACTCGGAGCTCTGTATTGGCCGATGCTTTGAACGCCATCAACAATGCCGAAAAGACCGGTAAGAGACAAGTT
TTAATTAGACCATCCTCCAAGGTTATTATTAAGTTCTTGCAAGTTATGCAAAAGCACGGTTACATTGGTGAATTTGA
ATACATTGACGACCATCGTTCCGGTAAGATCGTTGTTCAATTGAACGGTAGATTGAACAAGTGTGGTGTTATTTCTC
CAAGATTTAACGTTAAGATCGGTGACATTGAAAAATGGACCGCTAACTTGTTGCCAGCTAGACAATTCGGTTACGT
TATCTTAACCACCTCCGCCGGTATCATGGATCACGAAGAAGCTAGAAGAAAGCACGTCTCTGGTAAGATTTTGGGT
TTCGTCTACTAA

>RPS11A, YDR025W, SGDID:S000002432

ATGAGTACCGAGCTCACGGTTCAATCTGAAAGAGCTTTCCAAAAGCAACCACACATCTTCAACAACCCAAAAGTCA
AAACCTCCAAGAGAACCAAGAGATGGTACAAGAACGCTGGATTGGGTTTCAAAACCCCAAAGACCGCTATCGAGG
GTTCCTACATTGACAAGAAGTGTCCATTCACTGGTTTGGTCTCCATCAGAGGTAAGATTTTGACCGGTACTGTTGTT
TCCACCAAGATGCACAGAACTATTGTCATCAGACGTGCTTACTTACACTACATTCCAAAATACAACAGATACGAAA
AGAGGCATAAGAATGTTCCAGTCCACGTTTCCCCAGCTTTCAGAGTTCAAGTTGGCGACATCGTTACTGTCGGTCA
ATGTCGTCCAATCTCCAAGACCGTCCGTTTTAACGTTGTCAAGGTTTCCGCCGCTGCTGGTAAGGCTAACAAGCAA
TTCGCTAAGTTTTAA

>RPP1A, YDL081C, SGDID:S000002239

ATGAGCACCGAGTCTGCTTTGTCTTACGCTGCTTTGATCTTGGCTGATTCCGAAATTGAAATCTCCAGCGAAAAGTT
ATTGACTTTGACTAATGCTGCTAACGTCCCAGTCGAAAACATTTGGGCTGACATCTTCGCTAAGGCCTTAGATGGTC
AAAACTTGAAGGACTTGTTGGTCAATTTTTCTGCCGGTGCTGCTGCTCCAGCCGGTGTTGCCGGTGGTGTCGCCGG

TGGTGAAGCTGGTGAAGCTGAAGCCGAAAAAGAAGAAGAAGAAGCTAAGGAAGAATCTGACGATGACATGGGT
TTTGGCTTGTTCGAC

>RPP2B, YDR382W, SGDID:S000002790

ATGAAGTATTTGGCTGCCTACTTGTTGTTGGTCCAAGGTGGTAACGCCGCTCCATCTGCTGCTGATATCAAGGCTG
TTGTCGAATCTGTCGGCGCTGAAGTTGACGAAGCTAGAATCAACGAATTATTATCTTCCTTAGAAGGTAAGGGTTC
TTTAGAAGAAATCATCGCTGAAGGTCAAAAAAAGTTCGCCACTGTCCCAACTGGTGGTGCTTCTTCCGCTGCTGCT
GGTGCTGCTGGTGCCGCCGCTGGTGGTGACGCCGCCGAAGAAGAAAAGGAAGAAGAAGCTAAGGAAGAATCTG
ATGATGATATGGGTTTCGGTTTGTTTGAT

>RPL1A, YPL220W, SGDID:S000006141

ATGTCTAAAATTACCTCCTCCCAAGTTAGAGAACACGTTAAGGAATTGTTGAAGTACTCTAACGAAACTAAGAAGA
GAAACTTTTTGGAAACTGTTGAATTGCAAGTTGGTTTGAAGAACTACGATCCACAACGTGACAAGAGATTCTCCGG
TTCCTTGAAGTTGCCTAACTGTCCAAGACCAAACATGTCTATCTGTATCTTCGGTGACGCTTTCGACGTTGATAGAG
CTAAGTCCTGTGGTGTCGACGCTATGTCTGTTGACGATTTGAAGAAGTTGAACAAGAATAAGAAGTTGATCAAGA
AGTTGTCTAAGAAGTATAACGCTTTCATCGCTTCTGAAGTTTTGATTAAGCAAGTTCCAAGATTGTTGGGTCCACAA
TTGTCTAAGGCTGGTAAGTTCCCAACCCCTGTTTCTCACAACGACGATTTGTACGGTAAGGTTACCGACGTTAGATC
CACCATCAAGTTCCAATTGAAGAAGGTTTTGTGCTTGGCCGTCGCTGTTGGTAACGTCGAAATGGAAGAAGATGTT
CTAGTCAACCAAATCTTGATGTCCGTCAACTTCTTCGTCTCCCTATTAAAGAAGAATTGGCAAAACGTCGGTTCTTT
GGTTGTCAAGTCTTCTATGGGTCCAGCTTTCAGATTGTAT

>RPL2A, YFR031C-A, SGDID:S000002104

ATGGGACGTGTCATCAGAAACCAACGTAAGGGTGCCGGTTCCATTTTCACTTCTCATACTAGATTGCGTCAAGGTG
CCGCCAAGTTGAGAACCTTGGACTACGCCGAAAGACACGGTTACATTAGAGGTATTGTCAAGCAAATTGTTCACG
ACTCTGGTAGAGGTGCCCCATTGGCCAAGGTCGTTTTTAGAGATCCATACAAGTACAGATTGAGAGAAGAAATTTT
CATCGCCAACGAAGGTGTTCATACCGGTCAATTTATCTACGCTGGTAAGAAGGCCTCCTTGAACGTTGGTAACGTT
TTGCCATTGGGTTCCGTTCCAGAAGGTACTATCGTTTCTAACGTCGAAGAAAAGCCAGGTGATCGTGGTGCTTTGG
CTAGAGCCTCTGGTAACTACGTCATCATTATTGGTCACAACCCTGATGAAAATAAGACCAGAGTTCGTTTGCCATCT
GGTGCTAAGAAGGTCATTTCTTCTGACGCTAGAGGTGTTATTGGTGTTATCGCTGGTGGTGGTAGAGTCGATAAG
CCACTATTAAAGGCCGGTAGAGCCTTCCACAAGTATAGATTGAAGCGTAACTCTTGGCCAAAGACCAGAGGTGTC
GCTATGAACCCAGTCGACCACCCACACGGTGGTGGTAACCACCAACACATCGGTAAGGCCTCCACTATTTCTCGTG
GTGCCGTCTCTGGTCAAAAAGCTGGTTTAATCGCTGCTAGACGTACCGGTTTGTTGAGAGGTTCCCAAAAGACCCA
AGAC

>RPL3, YOR063W, SGDID:S000005589

ATGTCCCATCGGAAGTACGAAGCTCCAAGACATGGTCACTTGGGTTTCTTGCCAAGAAAGAGAGCTGCTTCCATTA
GAGCTAGAGTCAAGGCTTTCCCAAAGGACGACAGATCTAAGCCAGTCGCTTTGACCTCTTTCTTGGGTTACAAGGC
TGGTATGACCACTATTGTTAGAGACTTGGACAGACCAGGTTCCAAGTTCCACAAGAGAGAAGTTGTTGAAGCTGTT
ACCGTCGTTGATACCCCACCAGTTGTTGTTGTTGGTGTCGTCGGTTACGTTGAAACTCCAAGAGGTTTGAGATCCTT
GACTACTGTCTGGGCTGAACACCTAAGCGATGAAGTCAAGAGAAGATTCTACAAGAACTGGTACAAGTCCAAGAA
GAAGGCTTTCACTAAGTACTCCGCTAAGTACGCCCAAGACGGTGCTGGTATCGAAAGAGAATTGGCAAGAATTAA
GAAGTACGCTTCTGTCGTTCGTGTCTTGGTTCATACTCAAATTCGTAAGACCCCATTGGCTCAAAAGAAGGCTCACT
TAGCTGAAATTCAATTGAACGGTGGTTCTATTTCCGAAAAGGTCGACTGGGCTAGAGAACACTTTGAAAAGACCG
TTGCTGTTGATTCTGTTTTCGAACAAAACGAAATGATTGATGCCATCGCCGTTACTAAGGGTCACGGTTTCGAAGG

TGTCACCCATAGATGGGGTACCAAGAAGTTGCCAAGAAAGACTCACAGAGGTTTGAGAAAAGTCGCTTGTATCGG
TGCCTGGCACCCAGCTCATGTCATGTGGTCTGTTGCTAGAGCTGGTCAAAGAGGTTACCACTCTAGAACTTCTATC
AACCATAAGATCTACAGAGTCGGTAAGGGTGACGACGAAGCTAACGGTGCTACTTCTTTTGACAGAACCAAGAAG
ACCATCACTCCAATGGGTGGTTTCGTCCATTACGGTGAAATCAAGAACGACTTCATTATGGTTAAGGGTTGTATTCC
AGGTAACAGAAAAAGAATTGTCACCCTACGTAAGAGCTTGTACACTAACACCAGCAGAAAGGCTTTGGAAGAAGT
TTCTTTGAAGTGGATTGACACCGCTAGTAAGTTCGGTAAGGGTAGATTCCAAACCCCAGCTGAAAAGCACGCTTTC
ATGGGTACCTTGAAAAAGGATTTG

>RPL4A, YBR031W, SGDID:S000000235

ATGTCTAGACCTCAAGTTACTGTTCACTCCTTGACCGGTGAAGCTACCGCTAACGCTTTGCCATTGCCTGCTGTTTTC
TCCGCTCCAATTAGACCAGACATCGTTCACACTGTTTTCACTTCCGTTAACAAGAACAAGCGTCAAGCTTACGCTGT
CTCCGAAAAGGCCGGTCACCAAACTTCTGCTGAATCCTGGGGTACCGGTAGAGCCGTTGCCAGAATCCCAAGAGT
CGGTGGTGGTGGTACTGGTAGATCCGGTCAAGGTGCTTTCGGTAACATGTGTAGAGGTGGTAGAATGTTCGCTCC
AACTAAGACTTGGCGTAAGTGGAATGTTAAGGTTAACCACAACGAAAAGAGATACGCTACTGCTTCCGCTATCGCC
GCTACCGCCGTTGCTTCCTTGGTTTTGGCCAGAGGTCACCGTGTCGAAAAGATCCCAGAAATTCCATTGGTTGTTTC
TACTGACTTGGAATCCATCCAAAAGACTAAAGAAGCTGTCGCTGCTTTGAAGGCCGTCGGTGCTCACTCTGATCTA
TTGAAGGTTTTGAAGTCTAAGAAGTTGAGAGCTGGTAAGGGCAAGTACCGTAACCGTAGATGGACCCAACGTAG
AGGTCCATTGGTCGTTTACGCTGAAGATAACGGTATTGTCAAGGCTTTGCGTAACGTCCCAGGTGTTGAAACCGCT
AACGTCGCTTCCTTGAACTTGTTGCAATTGGCTCCAGGTGCTCATTTGGGCCGTTTTGTCATCTGGACTGAAGCCGC
TTTTACTAAGTTAGACCAAGTCTGGGGTTCTGAAACTGTCGCTTCCTCTAAGGTCGGTTACACCTTGCCATCCCACA
TTATTTCTACCTCCGACGTTACCAGAATCATCAACTCCTCTGAAATTCAATCCGCTATCAGACCAGCCGGTCAAGCC
ACTCAAAAGAGAACCCACGTGTTGAAGAAGAACCCATTGAAGAACAAGCAAGTCTTGTTGAGATTGAACCCATAC
GCTAAGGTTTTCGCCGCCGAAAAGTTGGGTTCTAAGAAGGCTGAAAAGACCGGTACTAAACCTGCTGCTGTTTTCA
CTGAAACCTTGAAGCACGAC

>RPL5, YPL131W, SGDID:S000006052

ATGGCCTTTCAGAAGGATGCTAAGTCTTCCGCTTACTCTTCCAGATTTCAAACCCCATTCAGAAGAAGAAGAGAAG
GTAAGACCGACTACTACCAAAGAAAGAGACTAGTTACTCAACACAAGGCTAAGTACAACACCCCAAAGTACCGTTT
GGTTGTTAGATTCACCAACAAGGACATCATTTGTCAAATCATCTCCTCTACCATTACCGGTGACGTTGTTTTGGCCG
CTGCTTACTCTCACGAATTGCCAAGATACGGTATCACTCACGGTTTGACTAACTGGGCTGCTGCTTACGCTACCGGT
TTATTAATCGCTAGAAGAACTTTGCAAAAATTGGGTTTGGATGAAACCTACAAGGGTGTTGAAGAAGTCGAAGGT
GAATACGAATTGACTGAAGCTGTTGAAGACGGTCCAAGACCATTCAAGGTTTTCTTAGACATCGGCTTGCAAAGAA
CTACTACCGGTGCTAGAGTTTTCGGTGCTTTGAAGGGTGCTTCTGATGGTGGTTTGTACGTCCCACACTCTGAAAA
CCGTTTCCCAGGTTGGGACTTCGAAACTGAAGAAATCGACCCTGAATTGTTAAGATCCTACATCTTCGGTGGTCAC
GTCTCTCAATACATGGAAGAATTGGCTGATGACGACGAAGAACGTTTCTCCGAATTGTTCAAGGGTTACTTGGCCG
ATGATATCGACGCTGATTCCTTGGAAGACATTTACACCTCCGCTCACGAAGCTATTAGAGCCGATCCTGCCTTCAAG
CCAACCGAAAAAAAGTTCACTAAGGAACAATACGCTGCCGAATCTAAAAAGTACAGACAAACTAAGCTATCTAAG
GAAGAAAGAGCTGCTAGAGTCGCTGCTAAGATCGCTGCTTTGGCTGGTCAACAA

>RPL6A, YML073C, SGDID:S000004538

ATGTCTGCTCAGAAGGCTCCAAAGTGGTACCCATCTGAAGACGTCGCTGCTTTGAAAAAGACTCGTAAGGCTGCC
AGACCACAAAAGTTGAGAGCTTCCTTGGTCCCAGGTACTGTTTTGATTTTGTTGGCTGGTAGATTTCGTGGTAAGC
GTGTCGTCTACCTAAAACACTTGGAAGACAACACCTTGTTGATCTCTGGTCCTTTCAAGGTTAACGGTGTCCCACTA
CGTCGCGTTAACGCTAGATATGTTATCGCTACTTCTACCAAGGTCTCTGTTGAAGGTGTTAATGTCGAAAAGTTTAA

CGTTGAATACTTCGCTAAGGAAAAGTTGACCAAGAAGGAAAAAAAGGAAGCTAACCTATTCCCAGAACAACAAAA
TAAGGAAATTAAGGCTGAAAGAGTCGAAGACCAAAAGGTCGTTGACAAGGCTTTGATCGCTGAAATTAAGAAGA
CTCCATTGCTAAAGCAATACTTATCTGCCTCTTTCAGTTTGAAGAATGGTGACAAACCACATATGTTAAAGTTC

>RPL7A, YGL076C, SGDID:S000003044

ATGGCTGCGGAGAAGATTTTGACTCCAGAATCCCAATTGAAGAAATCCAAGGCCCAACAAAAGACAGCTGAACAA
GTTGCTGCTGAAAGAGCTGCCAGAAAGGCCGCTAACAAGGAAAAGCGTGCTATCATCTTGGAAAGAAACGCTGCT
TACCAAAAGGAATACGAAACAGCTGAAAGAAACATTATCCAAGCTAAGAGAGACGCTAAGGCTGCTGGTTCTTAC
TATGTTGAAGCTCAGCACAAACTAGTTTTCGTCGTTCGTATTAAGGGTATCAACAAGATCCCACCAAAGCCTAGAA
AGGTCTTGCAATTGTTGAGACTAACTAGAATTAACTCGGGTACTTTCGTTAAGGTCACTAAAGCTACCTTGGAATT
GTTGAAGCTAATCGAACCATACGTCGCCTACGGTTACCCATCTTACTCCACCATCAGACAATTGGTTTACAAGAGA
GGTTTCGGTAAGATTAACAAGCAAAGAGTCCCATTGTCTGATAACGCCATCATCGAAGCTAACTTGGGTAAGTACG
GTATTTTGTCTATCGATGACTTGATCCACGAAATCATTACTGTCGGTCCACACTTCAAGCAAGCTAACAACTTCTTGT
GGCCATTTAAGTTGTCTAACCCATCCGGTGGTTGGGGTGTTCCACGTAAGTTCAAGCACTTCATCCAAGGTGGTTC
CTTCGGTAACAGAGAAGAATTTATTAACAAGTTGGTCAAGTCTATGAAT

>RPL8A, YHL033C, SGDID:S000001025

ATGGCTCCCGGGAAGAAGGTTGCTCCAGCTCCATTCGGTGCTAAGTCCACCAAATCCAACAAGACCAGAAACCCAT
TGACCCACTCCACCCCAAAGAACTTCGGTATTGGTCAAGCCGTTCAACCAAAGAGAAACTTGTCTAGATACGTTAA
GTGGCCAGAATACGTTAGAGTTCAAAGACAAAAGAAAATTTTGTCTATTAGATTGAAGGTCCCACCTACTATCGCT
CAATTTCAATACACCTTGGACAGAAACACTGCCGCTGAAACTTTCAAGTTGTTCAACAAGTACAGACCAGAAACCG
CCGCTGAAAAGAAGGAAAGATTGACTAAGGAAGCTGCTGCTGTCGCTGAAGGTAAGTCTAAACAAGACGCCTCCC
CAAAGCCATACGCTGTTAAGTACGGTTTGAACCACGTCGTCGCTTTGATTGAAAACAAGAAGGCTAAGTTGGTCTT
GATCGCTAACGACGTTGATCCAATCGAATTGGTCGTTTTCTTGCCAGCTTTGTGTAAGAAGATGGGTGTCCCATAC
GCTATTGTTAAGGGTAAGGCCAGATTGGGTACTTTGGTCAACCAAAAAACTTCCGCTGTCGCTGCCCTTACCGAAG
TTAGAGCTGAAGATGAAGCCGCTTTGGCCAAGTTGGTCTCAACTATCGACGCTAACTTCGCTGATAAGTACGACGA
AGTGAAGAAGCACTGGGGTGGTGGTATTTTGGGTAACAAGGCCCAAGCTAAGATGGACAAGAGAGCTAAGAACT
CTGACTCTGCT

>RPL9A, YGL147C, SGDID:S000003115

ATGAAGTATATTCAAACCGAACAACAAATTGAAGTTCCAGAAGGTGTTACCGTTTCTATTAAATCACGTATCGTTAA
GGTTGTCGGTCCAAGAGGTACCTTGACTAAAAACTTGAAGCACATCGACGTCACTTTTACTAAGGTTAACAACCAA
TTGATTAAGGTCGCTGTCCATAACGGTGGTAGAAAGCACGTCGCTGCTTTGAGAACTGTTAAGTCCTTGGTCGATA
ACATGATCACCGGCGTCACTAAGGGTTACAAGTACAAAATGAGATACGTTTACGCTCACTTTCCAATCAACGTCAA
CATCGTTGAAAAGGACGGTGCTAAGTTTATCGAAGTTAGAAACTTTTTGGGTGATAAGAAGATCCGTAACGTTCCA
GTCAGAGACGGTGTCACCATCGAATTCTCTACTAACGTCAAGGATGAAATCGTCTTGTCCGGTAACAGCGTTGAAG
ATGTTTCTCAAAACGCCGCGGACTTACAACAAATTTGTAGAGTTAGAAACAAGGACATCAGAAAGTTCTTGGATGG
TATTTACGTCTCCCACAAGGGTTTCATCACCGAAGACTTG

>RPL10, YLR075W, SGDID:S000004065

ATGGCGCGTCGGCCAGCTAGATGTTACAGATACCAAAAGAACAAGCCATACCCAAAGTCCAGATACAACAGAGCT
GTCCCTGACTCTAAGATTAGAATTTACGACTTGGGTAAGAAGAAGGCTACTGTCGACGAATTCCCATTGTGTGTCC
ATTTAGTCTCTAACGAATTGGAACAATTGTCCTCTGAAGCTTTGGAAGCTGCTAGAATCTGTGCTAACAAGTACAT
GACCACCGTCTCTGGTAGAGACGCCTTCCATTTGAGAGTTCGTGTTCACCCATTCCACGTTTTGAGAATTAACAAGA

TGTTGTCCTGTGCTGGTGCTGACAGATTGCAACAAGGTATGAGAGGTGCCTGGGGTAAGCCACATGGTTTGGCTG
CCAGAGTTGATATCGGTCAAATCATCTTCTCTGTTAGAACCAAGGATTCTAACAAGGACGTCGTCGTCGAAGGTTT
GCGTAGAGCTAGATACAAGTTTCCAGGTCAACAAAAGATCATCTTGTCCAAGAAGTGGGGTTTCACTAACTTGGAT
AGACCAGAATACTTGAAGAAGAGAGAAGCTGGTGAAGTCAAGGATGATGGTGCCTTCGTCAAGTTCTTGTCTAAG
AAGGGTAGCTTGGAAAACAACATTAGAGAATTCCCTGAATACTTCGCTGCTCAAGCT

>RPL11A, YPR102C, SGDID:S000006306

ATGTCCGCTAAGGCCCAAAACCCAATGAGAGACTTGAAGATTGAAAAGTTGGTTTTGAACATTTCCGTCGGTGAAT
CTGGTGACAGATTGACTAGAGCTTCTAAGGTCTTGGAACAATTGTCCGGTCAAACCCCAGTCCAATCTAAGGCTAG
ATATACAGTTAGAACCTTCGGTATTAGAAGAAACGAAAAGATCGCTGTCCACGTCACTGTTAGAGGTCCAAAGGCT
GAAGAAATTTTAGAAAGAGGTTTAAAAGTTAAGGAATACCAATTGAGAGATAGAAACTTCTCCGCCACTGGTAAC
TTCGGTTTCGGTATCGATGAGCACATCGATTTGGGTATTAAGTACGACCCTTCTATTGGTATCTTCGGTATGGACTT
CTACGTCGTTATGAACAGACCAGGTGCTAGAGTTACCAGACGTAAGAGATGTAAGGGTACTGTCGGTAACTCTCA
CAAAACCACCAAAGAAGACACTGTTTCTTGGTTCAAGCAAAAGTATGACGCCGACGTCTTAGATAAG

>RPL12A, YEL054C, SGDID:S000000780

ATGCCACCCAAATTCGACCCAAACGAAGTCAAGTACTTGTACTTGAGAGCTGTCGGTGGCGAAGTTGGTGCTTCTG
CTGCCTTAGCTCCTAAGATTGGTCCATTGGGTTTGTCTCCAAAGAAGGTCGGTGAAGACATTGCCAAAGCTACTAA
GGAATTCAAAGGTATTAAGGTCACTGTTCAATTGAAGATTCAAAATCGTCAAGCTGCTGCTTCCGTCGTTCCATCG
GCTAGTTCCCTAGTTATCACTGCCCTAAAGGAACCACCTAGAGATAGAAAGAAGGATAAGAATGTTAAACATTCCG
GCAATATCCAATTAGACGAAATCATCGAAATCGCTAGACAAATGAGAGATAAGTCTTTCGGTAGAACCTTAGCCTC
TGTTACTAAGGAAATTTTGGGTACCGCTCAATCCGTTGGTTGTAGAGTCGATTTTAAAAATCCACACGACATTATTG
AAGGTATCAACGCCGGTGAAATTGAAATTCCTGAAAAC

>RPL13A, YDL082W, SGDID:S000002240

ATGGCTATCTCTAAGAACTTGCCAATTTTGAAGAACCACTTCAGAAAGCACTGGCAAGAAAGAGTTAAGGTTCATT
TCGATCAAGCTGGTAAGAAGGTTTCTAGAAGAAACGCCAGAGCTACTAGAGCTGCTAAGATCGCTCCAAGACCAT
TGGACTTGTTACGTCCAGTCGTTAGAGCTCCTACCGTTAAGTACAATCGTAAGGTTAGAGCTGGTAGAGGTTTCAC
TTTAGCTGAGGTCAAGGCTGCCGGTTTAACCGCTGCTTATGCTAGAACCATTGGTATTGCTGTGGATCACCGTAGA
CAAAATAGAAATCAAGAAATTTTCGACGCTAATGTTCAAAGACTCAAAGAATATCAATCCAAAATCATCGTTTTTCC
TAGAAATGGCAAGGCTCCAGAAGCCGAACAAGTTTTATCTGCTGCCGCTACCTTCCCAATCGCTCAACCAGCTACT
GACGTTGAAGCCAGAGCTGTTCAAGATAACGGTGAATCTGCCTTCAGAACTCTAAGATTGGCTAGATCTGAAAAG
AAGTTCAGAGGTATTAGAGAAAAGAGAGCTAGAGAAAAAGCCGAAGCCGAAGCTGAAAAGAAGAAG

>RPL14A, YKL006W, SGDID:S000001489

ATGTCTACTGACTCCATCGTTAAGGCTTCTAACTGGAGATTGGTTGAAGTCGGTAGAGTTGTCTTGATTAAGAAGG
GTCAATCTGCTGGTAAGCTAGCTGCTATTGTTGAAATCATTGACCAAAAGAAAGTCTTAATTGACGGCCCAAAGGC
TGGTGTTCCAAGACAAGCTATTAACTTAGGTCAAGTCGTTTTGACCCCATTGACTTTTGCCTTGCCACGTGGTGCTC
GTACTGCTACTGTCTCCAAGAAGTGGGCTGCTGCCGCAGTTTGCGAAAAGTGGGCTGCTTCTTCATGGGCTAAAAA
GATCGCTCAAAGAGAAAGAAGAGCTGCTTTAACCGATTTCGAAAGATTTCAAGTGATGGTCTTGCGTAAGCAAAA
GCGTTACACCGTTAAGAAGGCTTTGGCTAAGGCT

>RPL15B, YMR121C, SGDID:S000004728

ATGGGTGCTTATAAGTACTTGGAAGAATTGGAAAGAAAGAAGCAATCCGACGTCTTAAGATTCTTGCAAAGGGTT
AGAGTTTGGGAATATAGACAAAAGAACGTCATCCATAGGGCTGCTAGACCCACCCGTCCTGATAAAGCTAGAAGA
TTGGGTTATAAGGCTAAACAAGGTTTCGTCATCTACAGAGTTAGAGTCAGAAGAGGAAATAGGAAAAGGCCAGTC
CCAAAAGGAGCTACTTACGGCAAGCCTACTAACCAAGGCGTTAACGAATTAAAGTACCAACGCTCGTTGAGAGCC
ACTGCTGAAGAAAGGGTCGGTAGAAGAGCTGCCAATTTGCGGGTTTTGAACTCTTACTGGGTTAATCAAGATTCC
ACCTACAAGTACTTCGAAGTCATCCTGGTCGATCCACAACATAAAGCCATCCGTAGAGACGCCAGATACAACTGGA
TTTGCAACCCAGTTCACAAACACCGTGAAGCCAGAGGTCTAACCGCTACTGGCAAAAAGTCTAGAGGCATAAACA
AAGGTCACAAGTTCAATAACACTAAGGCTGGTAGACGTAAGACTTGGAAGAGACAAAACACCTTATCCTTGTGGA
GATACAGAAAG

>RPL16A, YIL133C, SGDID:S000001395

ATGTCCGTCGAGCCTGTTGTTGTTATCGATGGTAAGGGTCATTTGGTTGGTAGATTGGCTTCTGTCGTTGCCAAGC
AATTGTTGAATGGTCAAAAGATTGTCGTTGTTAGAGCTGAAGAATTGAACATCTCTGGTGAATTCTTCAGAAATAA
GTTGAAATATCATGATTTCTTACGTAAGGCAACCGCTTTCAACAAAACTAGAGGTCCATTCCACTTTCGTGCTCCAT
CAAGAATTTTTTACAAGGCCTTGAGAGGTATGGTTTCCCATAAAACCGCTAGAGGTAAGGCTGCCTTAGAAAGATT
GAAAGTTTTCGAAGGTATCCCACCACCATACGATAAAAAGAAGAGAGTTGTTGTTCCACAAGCTTTACGTGTTTTA
CGTTTAAAGCCAGGTAGAAAGTACACTACCCTTGGTAAGTTATCTACCTCTGTTGGTTGGAAGTACGAAGATGTCG
TTGCTAAGTTAGAAGCTAAGAGAAAGGTTTCTTCTGCTGAATACTACGCTAAGAAACGTGCATTCACTAAAAAGGT
TGCTTCCGCTAACGCCACCGCCGCTGAATCTGATGTAGCTAAGCAATTGGCTGCCTTAGGTTAC

>RPL17A, YKL180W, SGDID:S000001663

ATGGCGCGTTATGGTGCTACCTCTACTAACCCAGCTAAGTCTGCCTCCGCTAGAGGTTCTTACTTGAGAGTTTCCTT
TAAGAACACTAGAGAAACTGCTCAAGCTATCAACGGTTGGGAATTGACTAAGGCTCAAAAGTACTTGGAACAAGT
TTTAGATCACCAAAGAGCTATCCCATTCCGTAGATTCAACTCCTCTATTGGTAGAACTGCTCAAGGTAAGGAATTCG
GTGTTACTAAGGCCAGATGGCCTGCTAAGTCCGTTAAGTTTGTCCAAGGTTTGTTGCAAAACGCTGCTGCTAACGC
TGAAGCTAAGGGTTTGGACGCCACCAAGTTGTACGTTTCCCACATTCAAGTCAACCAAGCCCCAAAACAAAGAAG
ACGTACTTACAGAGCTCATGGTAGAATTAACAAATACGAATCTTCTCCTTCTCACATCGAATTGGTCGTTACTGAAA
AGGAAGAAGCTGTTGCTAAGGCTGCCGAAAAGAAAGTTGTCAGATTGACCTCTAGACAAAGAGGTAGAATTGCT
GCTCAAAAAAGAATCGCTGCT

>RPL18B, YNL301C, SGDID:S000005245

ATGGGGATCGACCACACCTCCAAGCAACACAAGAGATCCGGTCACAGAACTGCTCCAAAGTCTGATAACGTTTACT
TAAAGTTGTTGGTTAAGTTGTATACTTTCTTGGCTAGAAGAACTGACGCTCCTTTCAACAAGGTTGTCTTGAAGGCG
TTGTTCTTGTCCAAGATCAACAGACCACCAGTGTCCGTTTCAAGAATCGCCAGAGCCTTGAAGCAAGAAGGTGCTG
CTAACAAAACTGTCGTTGTTGTCGGTACTGTCACCGATGATGCTAGAATCTTCGAATTCCCAAAAACTACCGTCGCC
GCTTTAAGATTCACTGCCGGTGCTCGTGCTAAGATTGTCAAGGCTGGTGGTGAATGTATCACCTTGGATCAATTGG
CTGTTAGAGCTCCAAAGGGTCAAAACACTCTAATCTTAAGAGGCCCACGTAACTCTAGAGAAGCTGTTAGACACTT
CGGTATGGGTCCTCATAAAGGTAAGGCTCCAAGAATTCTATCCACTGGTAGAAAGTTCGAAAGAGCTAGAGGTAG
AAGGAGATCCAAAGGTTTCAAGGTT

>RPL19A, YBR084C-A, SGDID:S000002156

ATGGCCAACTTGAGAACCCAAAAAAGATTGGCTGCCTCTGTTGTTGGTGTTGGTAAGAGAAAGGTCTGGTTGGAC
CCAAACGAAACTTCCGAAATCGCTCAAGCTAACAGTCGTAATGCTATCAGAAAGTTAGTCAAGAACGGTACCATTG
TTAAGAAGGCTGTCACCGTTCATTCTAAGTCTAGAACCAGAGCTCACGCTCAATCCAAGAGAGAAGGTAGACATTC

CGGTTACGGTAAGCGTAAGGGTACCAGAGAAGCTAGATTGCCATCTCAAGTAGTTTGGATCAGAAGATTGAGAGT
TTTAAGACGTTTGCTTGCTAAGTACAGAGATGCTGGTAAAATCGATAAGCACTTGTACCACGTCTTGTACAAGGAA
TCTAAGGGTAATGCTTTCAAGCACAAGAGAGCTTTAGTCGAACACATTATTCAAGCTAAAGCTGACGCTCAAAGAG
AAAAGGCCTTAAACGAAGAAGCTGAAGCTCGTCGTTTAAAGAACAGAGCCGCTAGAGATCGTAGAGCCCAAAGA
GTCGCCGAAAAGCGTGACGCCTTGTTAAAGGAAGATGCT

>RPL20A, YMR242C, SGDID:S000004855

ATGGCACATTTCAAGGAATATCAAGTCATCGGTAGAAGATTGCCAACCGAATCCGTTCCAGAACCTAAATTGTTTA
GAATGAGAATTTTCGCTTCCAACGAAGTCATCGCTAAGTCCAGATACTGGTACTTCTTGCAAAAGTTGCACAAGGT
CAAGAAGGCCTCCGGCGAAATCGTCTCTATTAATCAAATTAACGAAGCTCACCCTACTAAGGTTAAGAATTTCGGT
GTTTGGGTCAGATACGACTCTAGATCCGGTACCCACAACATGTACAAGGAAATTAGAGACGTTTCTAGAGTTGCTG
CTGTTGAAACTTTGTACCAAGACATGGCTGCTCGTCATCGTGCCAGATTCAGATCCATCCACATTTTGAAGGTCGCC
GAAATCGAAAAGACTGCTGACGTTAAAAGACAATACGTTAAGCAATTCTTGACTAAGGACTTGAAGTTTCCATTAC
CACACAGAGTTCAAAAGTCTACTAAGACCTTTTCTTACAAACGTCCATCAACCTTTTAC

>RPL21A, YBR191W, SGDID:S000000395

ATGGGCAAGTCTCACGGTTACAGATCTAGAACTAGATACATGTTTCAAAGAGACTTCAGAAAGCACGGTGCTGTTC
ACTTGTCAACCTACTTAAAAGTTTATAAGGTTGGTGACATCGTTGACATCAAGGCTAACGGTTCCATCCAAAAGGG
TATGCCTCATAAATTCTACCAAGGTAAGACTGGTGTTGTTTACAATGTTACTAAGTCTTCCGTCGGTGTTATCATCA
ATAAGATGGTTGGTAACCGTTACTTGGAAAAGAGATTGAACTTGCGTGTTGAACACATCAAGCATTCTAAATGTAG
ACAAGAATTCCTTGAACGTGTCAAGGCTAACGCCGCCAAGAGAGCTGAAGCCAAGGCTCAAGGTGTCGCTGTTCA
ACTAAAGCGTCAACCAGCCCAACCACGTGAATCTAGAATTGTCTCTACTGAAGGTAACGTTCCACAAACTTTAGCTC
CAGTCCCATACGAAACTTTTATC

>RPL22B, YFL034C-A, SGDID:S000006436

ATGGCCCCGAATACCTCTAGGAAACAAAAGGTAATAAAGACTTTGACTGTAGATGTCTCTTCCCCTACCGAAAACG
GTGTTTTCGACCCAGCCTCTTATAGTAAATATCTAATAGACCATATCAAGGTTGATGGGGCTGTCGGTAATTTGGG
TAACGCTATCGAAGTAACCGAAGACGGATCAATCGTTACAGTTGTCTCTTCCGCTAAGTTCTCCGGAAAGTATTTA
AAGTACTTGACTAAAAAATACCTAAAAAAAAACCAATTAAGAGATTGGATTAGATTTGTATCCATTAGACAAAACC
AGTATAAATTAGTCTTCTACCAAGTTACTCCAGAGGACGCCGATGAGGAAGAAGATGATGAA

>RPL23A, YBL087C, SGDID:S000000183

ATGTCTGGAAATGGTGCCCAAGGTACCAAGTTCAGAATTAGTTTGGGTTTGCCAGTTGGTGCTATTATGAACTGTG
CCGATAACTCCGGTGCTCGTAACTTGTACATCATCGCAGTTAAGGGTTCAGGCTCTAGACTAAATAGATTACCAGC
TGCTTCCTTGGGTGACATGGTCATGGCTACTGTCAAGAAGGGTAAGCCAGAACTAAGAAAAAAGGTCATGCCAGC
CATCGTCGTTAGACAAGCCAAGTCCTGGAGAAGAAGAGATGGTGTTTTTTTATACTTTGAAGACAACGCCGGTGTT
ATCGCCAACCCAAAGGGTGAAATGAAGGGTTCTGCTATCACTGGTCCAGTTGGTAAGGAATGTGCCGACTTGTGG
CCTAGAGTGGCCTCCAATTCTGGTGTCGTCGTT

>RPL24A, YGL031C, SGDID:S000002999

ATGAAAGTCGAGATCGACTCTTTCTCTGGCGCTAAGATTTACCCAGGTAGAGGTACTTTATTCGTTAGAGGTGATT
CTAAGATCTTCAGATTCCAAAACAGTAAGTCTGCTTCTTTGTTCAAGCAACGTAAAAACCCAAGAAGAATCGCTTG
GACCGTTTTGTTCAGAAAGCACCATAAAAAGGGTATCACTGAAGAAGTCGCTAAGAAGAGATCCAGAAAGACTGT
TAAGGCTCAACGTCCAATCACCGGTGCCTCTTTGGACTTGATCAAAGAAAGAAGATCCTTGAAGCCAGAAGTCAG

AAAGGCTAACAGAGAAGAAAAGTTGAAGGCTAACAAGGAAAAGAAAAAGGCTGAAAAGGCCGCCAGAAAAGCT
GAAAAAGCCAAGTCTGCTGGTACCCAATCCTCTAAGTTTTCTAAGCAACAAGCTAAGGGTGCTTTTCAAAAGGTTG
CTGCTACCTCAAGA

>RPL26A, YLR344W, SGDID:S000004336

ATGGCGAAGCAGTCCTTGGACGTTTCTTCCGATAGAAGAAAGGCTAGAAAAGCTTACTTTACCGCTCCATCATCCC
AAAGAAGAGTCTTGTTGTCCGCCCCATTGTCCAAGGAACTTCGTGCCCAATACGGTATTAAGGCTTTACCAATTCGT
CGTGACGACGAAGTTTTGGTTGTTAGAGGCTCTAAGAAGGGTCAAGAAGGTAAAATCTCCTCCGTTTATAGATTAA
AGTTCGCTGTTCAAGTTGACAAAGTCACTAAAGAGAAGGTTAACGGTGCCTCAGTTCCAATCAACTTGCATCCATC
TAAGTTGGTTATCACCAAGTTGCACTTGGATAAGGACAGAAAGGCTCTAATTCAAAGAAAGGGTGGTAAGTTAGA
A

>RPL27A, YHR010W, SGDID:S000001052

ATGGCCAAATTTTTAAAGGCCGGTAAGGTCGCTGTTGTTGTTAGAGGTAGATACGCTGGTAAGAAGGTCGTTATC
GTCAAGCCTCACGATGAAGGTTCTAAGTCCCATCCATTCGGTCATGCTTTAGTCGCTGGTATCGAACGTTACCCATT
GAAGGTTACTAAGAAGCACGGTGCTAAGAAAGTCGCTAAAAGAACTAAGATCAAGCCATTCATTAAGGTTGTTAA
CTACAACCACTTGTTGCCTACTCGTTACACCTTGGATGTTGAAGCCTTCAAGTCTGTCGTCTCCACCGAAACCTTCG
AACAACCATCTCAAAGAGAAGAAGCTAAGAAGGTTGTTAAAAAGGCTTTTGAAGAACGTCACCAAGCTGGTAAAA
ACCAATGGTTTTTCTCCAAATTGAGATTC

>RPL29, YFR032C-A, SGDID:S000006437

ATGGCAAAAAGCAAAAACCACACTGCTCATAACCAAACCAGAAAGGCTCACAGAAACGGCATCAAGAAGCCAAA
GACCTACAAGTACCCATCTCTAAAGGGTGTTGATCCTAAATTTAGAAGAAACCACAAGCACGCTTTGCATGGTACC
GCCAAGGCCTTGGCTGCTGCTAAGAAG

>RPL31B, YLR406C, SGDID:S000004398

ATGGCTGGGTTGAAGGATGTTGTTACCAGAGAATACACTATCAACCTACACAAGCGTTTGCACGGTGTTTCCTTCA
AGAAACGTGCCCCAAGAGCCGTTAAGGAAATCAAAAAGTTTGCTAAGTTGCATATGGGTACCGAAGATGTTAGAT
TGGCTCCAGAACTAAACCAAGCCATCTGGAAGAGAGGTGTCAAAGGTGTGGAATACAGACTAAGATTGAGAATC
AGTAGAAAAAGAAACGAAGAAGAAGATGCTAAAAACCCATTATTCTCTTACGTTGAACCAGTCTTAGTCGCTTCCG
CCAAAGGTTTGCAAACTGTCGTTGTCGAAGAAGACGCC

>RPL33B, YOR234C, SGDID:S000005760

ATGGCCGAGTCTCACCGTTTGTACGTTAAGGGTAAACATTTGTCTTACCAAAGATCTAAGAGAGTCAACAACCCAA
ACGTTTCCCTAATTAAGATCGAAGGTGTCGCTACTCCACAAGAAGCCCAATTTTACTTGGGTAAGAGAATTGCTTA
CGTCTATAGAGCCTCTAAGGAAGTTAGAGGTTCTAAAATTAGAGTCATGTGGGGTAAGGTTACTAGAACCCACGG
CAACTCCGGTGTTGTCAGAGCCACTTTCAGAAATAACTTGCCAGCTAAGACCTTCGGTGCTTCCGTTAGAATTTTCT
TATACCCATCCAACATT

>RPL34A, YER056C-A, SGDID:S000002135

ATGGCTCAGAGAGTTACCTTCAGAAGAAGAAACCCATACAACACCAGATCCAATAAGATTAAGGTCGTTAAGACT
CCAGGTGGTATCTTGAGAGCTCAACACGTTAAGAAATTGGCCACTAGACCAAAGTGTGGTGACTGTGGTTCTGCTT
TACAAGGTATTTCCACCTTGCGTCCACGTCAATACGCTACCGTCTCTAAGACTCATAAAACCGTTTCTCGTGCCTAC

GGTGGTAGCAGATGTGCTAACTGTGTTAAGGAAAGAATCATCAGAGCCTTCTTGATTGAAGAACAAAAGATTGTT
AAAAAGGTCGTTAAAGAACAAACTGAAGCCGCTAAGAAGTCCGAAAAGAAGGCCAAGAAG

>RPL35A, YDL191W, SGDID:S000002350

ATGGCTGGCGTCAAGGCTTACGAATTGAGAACCAAGTCTAAGGAGCAATTGGCCTCCCAATTGGTCGACTTGAAG
AAGGAATTGGCCGAATTAAAGGTTCAAAAGCTAAGTAGACCATCCTTGCCAAAGATCAAAACCGTTAGAAAATCC
ATCGCTTGTGTTTTGACCGTTATTAACGAACAACAAAGAGAAGCTGTCAGACAATTGTACAAAGGTAAGAAGTACC
AACCAAAGGACTTGAGAGCTAAGAAGACCAGAGCTTTGAGAAGAGCCTTGACCAAGTTCGAAGCTTCTCAAGTCA
CCGAAAAGCAAAGAAAGAAGCAAATCGCTTTCCCACAAAGAAAATACGCTATCAAGGCT

>RPL36A, YMR194W, SGDID:S000004807

ATGACTGTCAAAACCGGTATCGCCATCGGTTTGAACAAGGGAAAAAAGGTCACCTCTATGACCCCAGCTCCAAAG
ATTTCTTACAAGAAGGGTGCTGCTTCTAACCGTACAAAATTCGTCAGATCCTTGGTTAGAGAAATTGCTGGTCTATC
TCCATACGAACGTAGATTAATAGATTTGATTAGAAACTCTGGTGAAAAGAGAGCTAGAAAGGTCGCTAAGAAGAG
ATTGGGTTCTTTCACTAGAGCTAAAGCTAAGGTAGAAGAAATGAACAACATTATCGCCGCTAGCAGAAGACAC

>RPL37A, YLR185W, SGDID:S000004175

ATGGGAAAAGGCACTCCATCTTTCGGTAAGAGACACAACAAGTCTCACACCTTGTGTAACCGTTGTGGTAGAAGAT
CCTTCCACGTTCAAAAGAAGACTTGTTCTTCTTGTGGTTACCCTGCTGCTAAAACTAGATCCTACAACTGGGGTGCT
AAGGCTAAGCGTAGACATACTACCGGTACCGGTCGTATGAGATATTTGAAGCACGTTTCTAGAAGATTCAAGAAC
GGTTTCCAAACTGGTTCCGCTTCAAAGGCCTCAGCT

>RPL38, YLR325C, SGDID:S000004317

ATGGCGCGTGAGATCACTGATATTAAGCAATTCTTGGAATTGACTAGAAGAGCCGACGTTAAGACCGCTACCGTT
AAGATCAACAAGAAGTTGAACAAAGCTGGTAAACCATTCAGACAAACTAAGTTTAAGGTTAGAGGTTCTTCTTCTT
TGTACACCTTAGTTATCAACGACGCCGGCAAAGCTAAAAAGTTGATCCAATCTTTGCCACCAACCTTGAAGGTTAA
CAGATTG

>RPL40A, YIL148W, SGDID:S000001410

ATGCAGATCTTCGTTAAAACCTTGACCGGTAAAACTATTACCTTGGAAGTCGAATCTTCCGACACCATCGACAACGT
TAAATCTAAGATTCAAGACAAGGAAGGTATTCCACCAGACCAACAAAGATTGATTTTTGCTGGTAAGCAATTGGAA
GACGGTAGAACTTTGTCTGACTACAACATCCAAAAGGAATCTACTTTGCACTTGGTCTTGAGATTGAGAGGTGGTA
TTATCGAACCTTCCTTGAAGGCTTTGGCTTCTAAGTATAACTGTGACAAGTCAGTCTGTAGAAAGTGTTACGCTAGA
TTACCACCACGTGCTACTAATTGTCGTAAGAGAAAATGCGGTCACACTAACCAATTAAGACCAAAGAAGAAGTTGA
AG

>RPL41A, YDL184C, SGDID:S000002343

ATGCGTGCTAAATGGAGAAAGAAGAGAACTAGAAGACTTAAGAGAAAGAGAAGAAAGGTGCGGGCCAGATCCA
AG

>RPL42A, YNL162W, SGDID:S000005106

ATGGTCAATGTACCTAAGACCCGTAAGACCTACTGCAAAGGTAAGACTTGTAGAAAACACACCCAACACAAAGTT
ACCCAATATAAGGCTGGTAAAGCCTCTTTGTTCGCTCAAGGTAAGAGAAGATACGACAGAAAGCAATCCGGTTTC
GGTGGTCAAACTAAGCCAGTTTTCCACAAGAAGGCTAAGACCACCAAGAAGGTTGTTTTGCGTTTGGAATGTGTTA

AGTGTAAGACTAGAGCTCAATTGACCTTGAAGCGTTGTAAGCACTTCGAATTGGGTGGTGAAAAGAAGCAAAAGG
GTCAAGCCTTGCAATTT

>RPL43A, YPR043W, SGDID:S000006247

ATGGCAAAGCGTACCAAGAAGGTCGGTATCACCGGTAAGTACGGTGTCAGATACGGTTCTTCCTTGAGAAGACAA
GTTAAGAAGTTGGAAATCCAACAACACGCTAGATATGATTGTTCCTTCTGTGGTAAAAAGACTGTTAAGAGAGGT
GCTGCTGGTATCTGGACCTGTTCTTGTTGTAAGAAGACTGTCGCCGGTGGTGCTTACACCGTTTCTACTGCTGCTGC
TGCTACTGTCAGATCCACTATTAGACGTTTGAGAGAAATGGTCGAAGCC
